# Systematic temporal analysis of peripheral blood transcriptomes using *TrendCatcher* identifies early and persistent neutrophil activation as a hallmark of severe COVID-19

**DOI:** 10.1101/2021.05.04.442617

**Authors:** Xinge Wang, Mark Sanborn, Yang Dai, Jalees Rehman

**Author notes:** Please address correspondence to Jalees Rehman, The University of Illinois at Chicago, College of Medicine (M/C 868) 835 S. Wolcott Ave. Rm. E403, Chicago, IL 60612, USA.

## Abstract

Studying temporal gene expression shifts during disease progression provides important insights into the biological mechanisms that distinguish adaptive and maladaptive responses. Existing tools for the analysis of time course transcriptomic data are not designed to optimally identify distinct temporal patterns when analyzing dynamic differentially expressed genes (DDEGs). Moreover, there is a lack of methods to assess and visualize the temporal progression of biological pathways mapped from time course transcriptomic datasets. In this study, we developed an open-source R package *TrendCatcher* (https://github.com/jaleesr/TrendCatcher), which applies the smoothing spline ANOVA model and break point searching strategy to identify and visualize distinct dynamic transcriptional gene signatures and biological processes from longitudinal datasets. We used *TrendCatcher* to perform a systematic temporal analysis of COVID-19 peripheral blood transcriptomes, including bulk RNA-seq and scRNA-seq time course data. *TrendCatcher* uncovered the early and persistent activation of neutrophils and coagulation pathways as well as impaired type I interferon (IFN-I) signaling in circulating cells as a hallmark of patients who progressed to severe COVID-19, whereas no such patterns were identified in individuals receiving SARS-CoV-2 vaccinations or patients with mild COVID-19. These results underscore the importance of systematic temporal analysis to identify early biomarkers and possible pathogenic therapeutic targets.

## Introduction

Time-course transcriptomic profiling has been widely used to study and model dynamic biological processes in cells (1). By profiling mRNA levels during consecutive time points, researchers can infer dynamic responses to various external cues that cannot be observed by looking at only initial and terminal states. Recent improvements in high-throughput RNA sequencing (RNA-seq) technologies, including single-cell RNA-sequencing (scRNA-seq) provide viable approaches to study dynamic gene expression changes (2). Especially scRNA-seq allows for the in-depth analysis of temporal changes in distinct cell populations, thus providing insights into the heterogeneity and dynamics of responses to environmental cues or pathogenic stimuli. However, the analysis and visualization of longitudinal bulk RNA-seq data or scRNA-seq can be computationally challenging.

Currently, there are two predominant strategies for the analysis of sequential transcriptomic datasets. One strategy treats the sampling time points as categorical variables and is based on generalized linear models (GLMs). GLM-based packages include DESeq2 (3), edgeR (4) and limma (5). A complementary strategy is to treat time as a continuous variable and fit the time expression data into a spline-like model. These methods include DESeq2Spline (DESeq2 adopted with spline model for temporal RNA-seq datasets) fitting, ImpulseDE2 (6) and Next maSigPro (7). The former strategies focus on the magnitude of change instead of the time order of gene expression, and may also suffer from a relative loss of statistical testing power, especially if many time points are assessed (6). The latter strategy increases the power of detecting dynamic genes, but is based on strong model assumptions which are not optimally suited for multiphasic responses, such as gene trajectory patterns that reflect initial acute stimuli followed by counter-regulatory compensatory responses. Furthermore, there is a lack of tools that leverage existing knowledge of functional pathway databases to infer and visualize pathway trajectories instead of individual gene trajectories only.

We found this is challenging in a complex disease, such as coronavirus disease 19 (COVID-19) caused by the SARS-CoV-2 virus infection (8). COVID-19 is characterized by distinct disease progression patterns that suggest diverse host immune responses (9). Patients with severe disease exhibit profound inflammatory responses and immunopathology (10). COVID- 19 immunophenotyping studies involve a large number of time points from corresponding RNA-seq and scRNA seq datasets (11, 12). Existing approaches for temporal transcriptomic analysis either do not take the temporal sequence into account when identifying dynamic differentially expressed genes (DDEGs) using pairwise comparison between time points, nor systematically analyze pathways that are dysregulated at defined time points.

In this study, we developed *TrendCatcher*, an open-source R package tailored for longitudinal bulk RNA-seq and scRNA-seq analysis. *TrendCatcher* uses a framework that combines the smooth spline ANOVA model and break point searching strategy, which identifies inflection points when gene expression trends reverse. We show that *TrendCatcher* outperformed commonly used methods for longitudinal RNA-seq analysis when using simulated time course data for benchmarking. We also analyzed bulk RNA-seq and scRNA-seq gene expression profiles of peripheral blood cells in COVID-19 patients at various disease time points. *TrendCatcher* allowed us to identify and visualize dynamic gene expression signature profiles in peripheral blood that were associated with poor disease outcomes during the early phases of disease, and could thus serve as novel mechanistic targets as well as early biomarkers for patient prognostication.

## Results

### *TrendCatcher* accurately identifies DDEGs in simulated datasets

First, we tested the prediction performance of the *TrendCatcher* platform (**Figure 1A**) using a set of simulated time course RNA-seq datasets, because simulated data provides defined standards to assess the accuracy of novel analytical platforms. We considered a comprehensive collection of gene temporal trajectory patterns to simulate a set of realistic data with biological characteristics (1). We embedded 10,000 simulated trajectories with varied temporal patterns, including 90% non-dynamic trajectories, 2.5% monotonous transition trajectories (continuously increasing or continuously decreasing gene expression levels throughout the time course), 2.5% impulse shaped single break point trajectories (only one temporal inflection point, i.e. up-peak-down or down-trough-up), 2.5% two break point trajectories and 2.5% three break point trajectories (multimodal dynamic response, e.g., a combination of 2 or more basic types of trajectories). Compared to DESeq2, DESeq2Spline and ImpulseDE2, *TrendCatcher* had the highest area under the ROC curve (AUC) in a mixed simulated dataset for time-course data with 7 time points (**Figure 1B**). We also tested each model’s performance on a varying number of time points, including 3, 5, 7, 9 and 11 time points. As shown in **Figure 1C**, *TrendCatcher* had the highest prediction AUC across all time points, with the AUC values range from 0.88 to 0.90.

**Figure 1.**
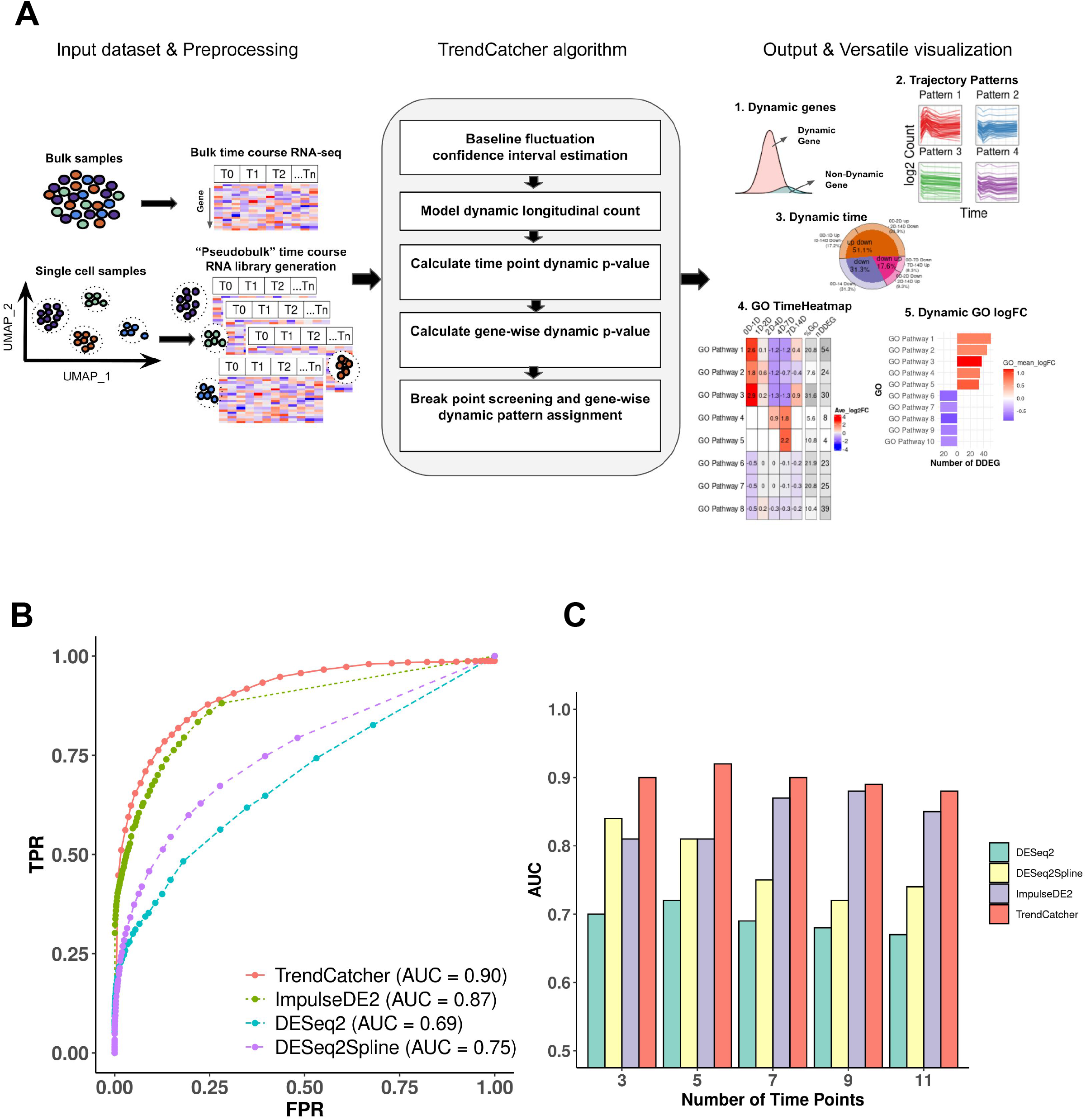
Overview and benchmarking of of *TrendCatcher*. (A) *TrendCatcher* framework. *TrendCatcher* preprocess input data, which includes creating cell-type specific “pseudobulk” datasets for temporal analysis when single-cell data is used. *TrendCatcher’*s core algorithm is composed of five main steps. *TrendCatcher*’s output includes four main types of visualizations and DDEG identification (numbered 1-5). (B) *TrendCatcher* prediction ROC for a 7 time point simulated dataset compared to DESeq2, DESeq2Spline and ImpulseDE2, with mixed trajectories. (C) *TrendCatcher* prediction performance (AUC) across different number of time points, from 3 to 11 time points. *TrendCatcher’*s AUC values across time points from 3 to 11 are, 0.90, 0.92, 0.90, 0.89 and 0.88.

We next evaluated the prediction performance for each type of temporal trajectories. As shown in (**Supplemental Figure 1**), although other three methods achieved slightly higher accuracy than *TrendCatcher* in monotonic trajectories, their AUCs dropped markedly when more complicated trajectories were embedded. DESeq2 only achieved AUC values of 0.49 to 0.62 for both bi-phasic trajectory and multimodal trajectory. The DESeq2Spline approach using a spline curve fitting model also dropped to an AUC of approximately 0.7 once multiphasic trajectories were introduced. These results suggests that existing approaches for longitudinal or time course analyses are well-suited for monotonic trajectories (continuously up or continuously down) but that *TrendCatcher* maybe more broadly applicable, because it identifies monotonic, biphasic (up-down, down-up) and multi-phasic shifts in gene expression, which are especially important in complex pathological setting when initial biological responses are followed by counter-regulatory adaptive or maladaptive responses.

### *TrendCatcher* identifies rapid but transient upregulation of interferon signaling in peripheral blood following SARS-CoV-2 infection in a non-human primate model

To define the key dynamic gene signatures associated with SARS-CoV-2 infection in peripheral blood, we first analyzed the global transcriptomics profiles from a non-human primate dataset (13), in which samples were collected on Days 0 (uninfected controls), 1, 2, 4, 7, 10 and 14 following the live SARS-CoV-2 inoculation. This experiment in non-human primates had the advantage of clearly defining the timing of inoculation and following the time course of dynamic genes. *TrendCatcher* identified 962 DDEGs out of 12,754 total genes, accounting for 7.6% of total expression, suggesting that over 90% of the expressed genes in the peripheral blood remain close to the baseline expression levels even in the setting of the SARS-CoV-2 infection. We observed two major types of dynamic trajectories: (a) 167 genes followed a biphasic “0D-2D up, 2D-14D down” pattern, with their expression level peaking at Day 2 and gradually returning close to baseline levels at Day 14 (**Figure 2A**). These dynamic genes were primarily associated with host defense biological pathways, such as defense response to viruses, regulation of viral life cycle and type I interferon signaling pathways (**Figure 2B**). (b) 263 genes follow a “0D-14D down” pattern (**Figure 2A**). This set of genes followed a monotonous trajectory, with their expression gradually decreasing until Day 14. Interestingly, we found these genes were primarily associated with mitochondrial ATP synthesis and oxidative phosphorylation, suggesting that disruption of mitochondrial respiration and ATP generation may be an important feature of the circulating cells transcriptomic shift in a SARS-CoV-2 infection (**Figure 2C**). *TrendCatcher* assigned trajectory pattern types to all dynamic genes, and provided hierarchical pie charts to visualize the composition of trajectory patterns (**Supplemental Figure 2A**).

**Figure 2.**
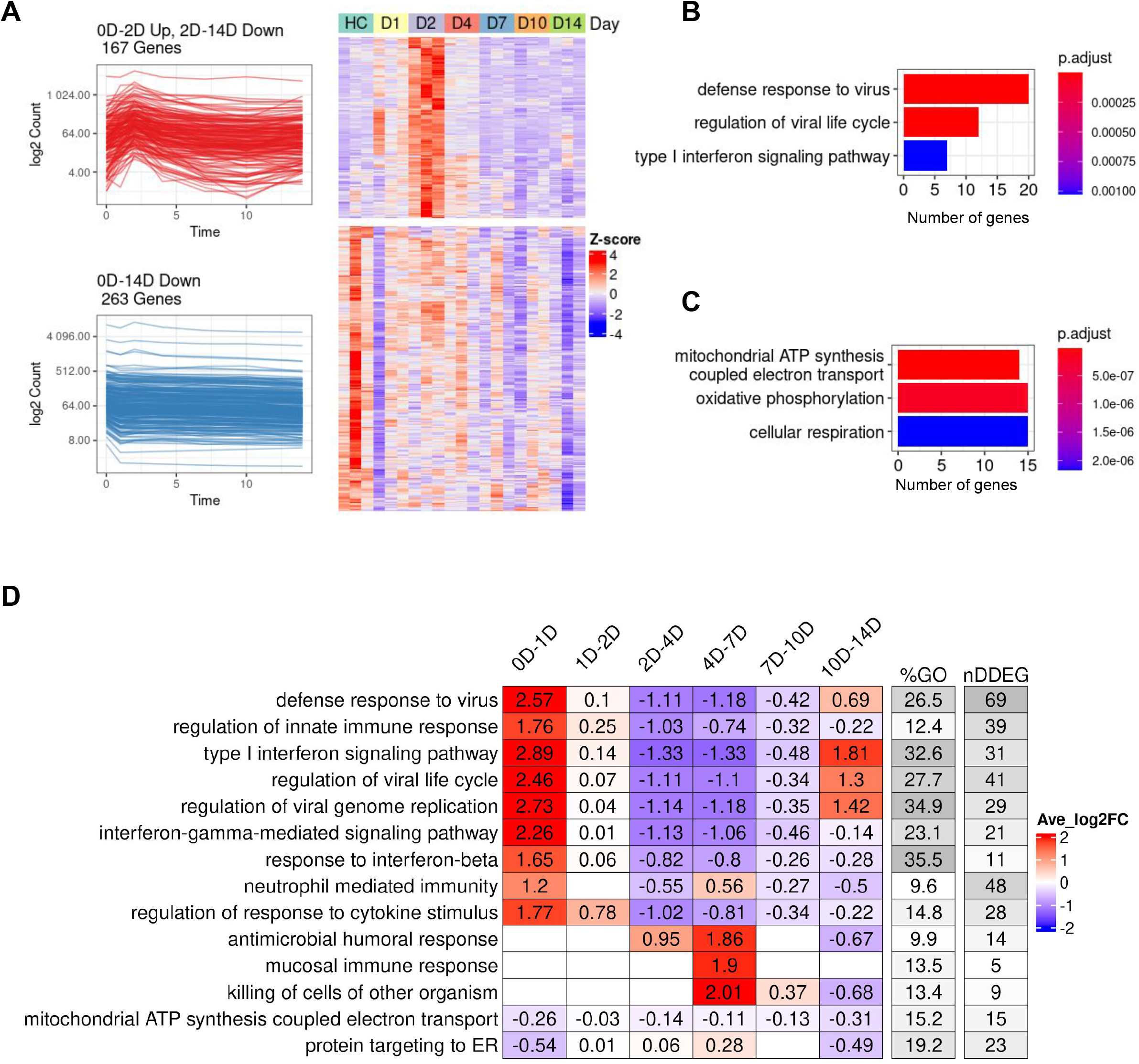
Dynamic gene expression in peripheral blood following SARS-CoV-2 inoculation in a non-human primate model. (A) Analysis of the two predominant trajectory patterns in the non-human primate peripheral blood RNA-seq data from Day 0 to Day 14. The top-left figure represents 167 DDEGs following an up-down expression pattern, which peaked at Day 2 then slowly decreased until Day 14. The top-right figure represents their expression using a traditional z-score normalized heatmap. The bottom-left figure represents 263 DDEGs following a monotonic down-regulated trajectory pattern and their gene expression values were represented in the corresponding heatmap on the right. (B) Top 3 Gene ontology (GO) enrichment analysis pathways using 167 DDEGs from trajectory pattern “0D-2D Up, 2D-14D Down”. (C) Top 3 Gene ontology (GO) enrichment analysis pathways using 263 DDEGs from trajectory pattern “0D-14D Down”. (D) *TimeHeatmap* of the top 15 dynamic pathways and their dynamic time windows visualizes the temporal patterns. Each column represents a time window. The “%GO” column represents the percentage of DDEGs found in the corresponding pathway. The “nDDEG” column represents number of DDEGs found in the corresponding pathway. Color represents the averaged log fold-change of the DDEGs within each time window for the corresponding pathway.

To systematically assess and visualize the dynamic programming of the top biological pathways associated with SARS-CoV-2 infection, *TrendCatcher* generated a *TimeHeatmap*.

The *TimeHeatmap* function of *TrendCatcher* visualizes shifts in pathways by displaying the mean-fold change of individual DDEGs in a given pathway at defined time points while also depicting the number of dynamic genes and the percentage of dynamic genes within that pathway (**Figure 2D**). This quantifies the pathway level shifts over time, and allows for the magnitude of pathway change to be visualized. Together with the number and fraction of dynamic genes in a given pathway, to gauge the relative importance of the pathway during the time course.

During initial infection (Day 0 to Day2), pathways related to innate immune response and interferon pathways were highly upregulated. Examples are upregulation of pathways such as defense response to virus and regulation of innate immune response increased with an average log2 fold-change of 2.57 and 1.76 within the first day. Mucosal immune response, antimicrobial humoral response and killing of cells of other organisms were activated during the later stage of infection (Day 4 to Day 7), with an average log2 fold-change around 2. On the other hand, mitochondrial ATP synthesis coupled electron transport and protein targeting to ER, on the other hand, were gradually down-regulated until Day 14. Dynamic gene signatures from type I interferon signaling pathway and mitochondrial ATP synthesis coupled electron transport were shown using traditional heatmaps in **Supplemental Figure 2B,C**. The temporal analysis of this non-human primate dataset highlights the rapidity of interferon signaling activation btu also underscores that this initial burst of immune activation is transient and followed by a gradual downregulation of the antiviral interferon responses during the first week following infection.

### Increased generation of immunoglobulin synthesis in plasma B cells as early as Day 1 of symptom onset in patients diagnosed with SARS-CoV-2 infection

Next, we analyzed the longitudinal gene expression profiles of peripheral blood mononuclear cells (PBMCs) obtained from patients diagnosed with a SARS-CoV-2 infection who were admitted to the hospital, but predominantly had uncomplicated disease progression with 4 out 5 patients showing only mild symptoms (14) (**Figure 3A**). The study performed single cell RNA-seq analysis on PBMCs, thus allowing for a cell-type specific analysis of gene expression shifts in distinct B cell subtypes, T cell subtypes and NK cells. We adopted the cell-type labels from the original study and generated pseudo-bulk RNA-seq datasets for each cell type in order to quantify changes in DDEGs for a specific cell type. We also adopted the time annotation using stages, by binning the disease processes of COVID-19 patients from symptom onset to discharge into stages 0, 1, 2, 3 and 4. We defined Stage 0 as the time point of samples obtained from healthy controls and stages 1, 2, 3 and 4 represented Day 1 to Day 16 from symptom onset. We only applied *TrendCatcher* to cell types containing more than 1,000 cells at any given stage to ensure the robustness of the results.

**Figure 3.**
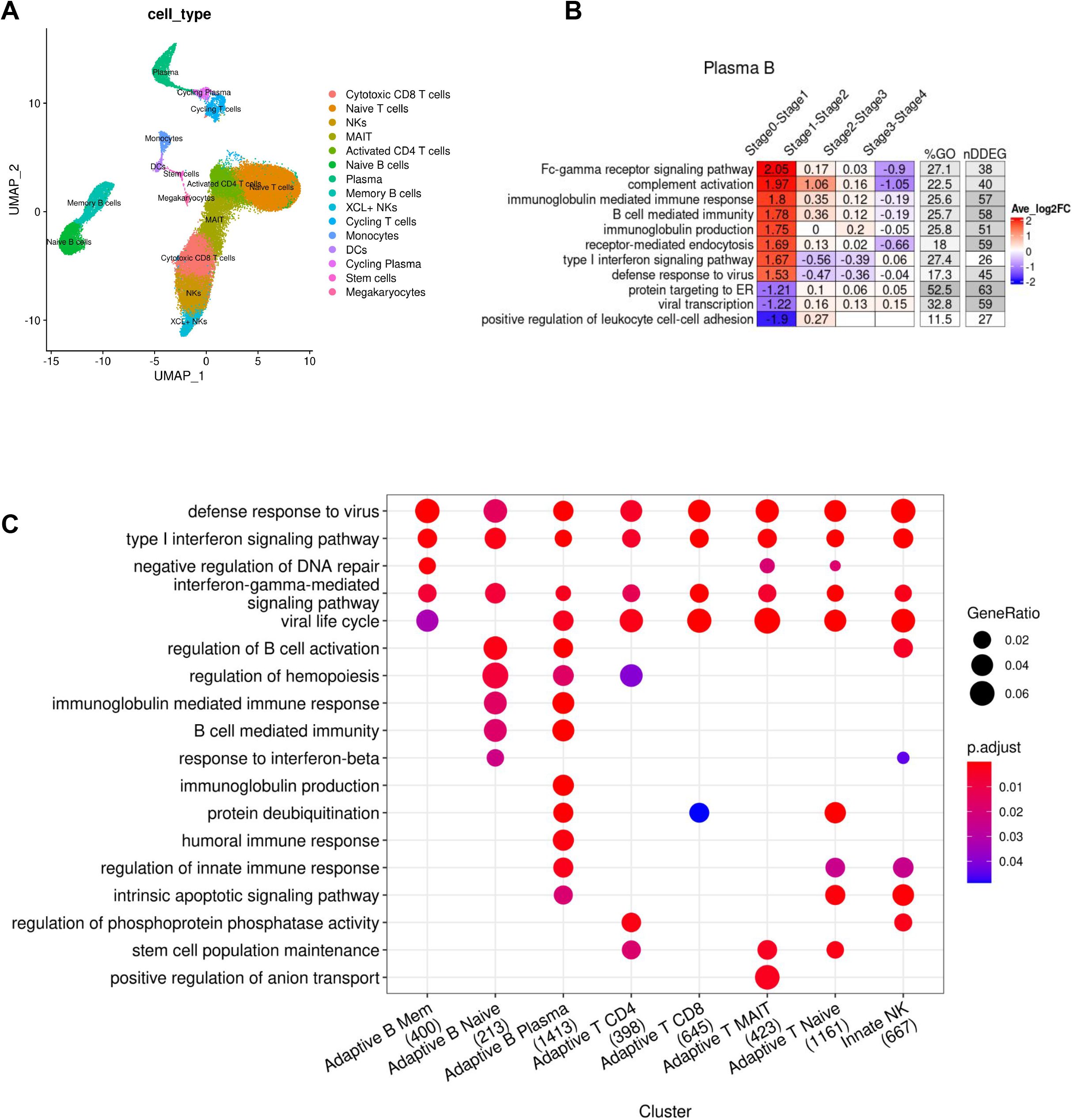
Cell-type specific dynamic gene expression in peripheral blood mononuclear cells following SARS-CoV-2 infection in patients. (A) UMAP visualization of single cell RNA-seq PBMC dataset (Zhu et al.) with annotated cell types from the original study. (B) *TimeHeatmap* of top dynamic biological pathway from plasma B cells. Each column represents a time window (stage). Stage 0 represents uninfected baseline. Each row represents the most dynamic biological pathways. Color represents the averaged log fold-change of the DDEGs within each time window from the corresponding pathway. (C) Top GO enrichment comparison analysis using DDEGs from each cell type. Dot size represents gene ratio. Dot color represents GO enrichment p-value. The number in the x-axis labels represent the number of DDEGs for each cell type.

*TrendCatcher* identified 400 DDEGs in memory B cells, 213 DDEGs in naive B cells, 1,413 DDEGs in plasma B cells, 398 DDEGs in CD4^+^ T cells, 645 DDEGs in CD8^+^ T cells, 423 DDEGs in MAIT, 1,161 DDEGs in naive T cells and 667 DDEGs in NK cells. The *TimeHeatmap* of plasma B cells visualized the most dynamic biological pathways. Importantly, this temporal analysis of single cell RNA-seq data showed how rapidly plasma B cells ramp up the upregulation Fc-gamma receptor signaling and immunoglobulin synthesis as early as stage 1 (which corresponds to Day 1 of symptom onset) (**Figure 3B**). However, not all immunoglobulin genes are upregulated during the same temporal phase. As shown in (**Supplemental Figure 3**), genes involved in immunoglobulin synthesis show distinct temporal patterns. Due to the comparatively lower number of plasma B cells, the prominence of such increased immunoglobulin changes may be diluted in peripheral blood bulk RNA-seq analysis or PBMC bulk RNA-seq analysis. However, temporal analysis of all peripheral blood mononuclear cell types, across the whole time course demonstrated increases in type I interferon signaling and defense responses to viruses as the most prominent changes over time (**Figure 3C**), thus mirroring the responses to SARS-CoV-2 we observed in the peripheral blood of non-human primates (**Figure 2**). To define cell-type specific temporal dynamics, we generated *TimeHeatmaps* for individual PBMC cell types and found significant upregulation of Type I interferon signaling in T cells, NK cells and memory B cells during Stage 1 but subsequent downregulation by Stage 2 of the disease in patients with mild symptoms (**Supplemental Figure 4A-C, Supplemental Figure 5A-D**).

### *TrendCatcher* identifies early and persistent neutrophil activation as a hallmark of severe COVID-19

We then assessed whether a systematic temporal analysis of gene expression trends could be used to distinguish disease severity and prognosis of COVID-19 patients. We thus applied *TrendCatcher* to a time course human whole blood bulk RNA-seq dataset (11), which contained longitudinal whole blood transcriptomes from COVID-19 patients with mild, moderate and severe clinical outcomes. *TrendCatcher* identified 77 DDEGs from the mild group, 226 DDEGs from the moderate group, and 1,205 DDEGs from the severe group. Only 42 DDEGs were shared among these three groups (**Figure 4A**), and these were primarily associated with B cell mediated immunity, Fc-gamma receptor signaling pathways, humoral immune response and lymphocyte mediated immunity (**Figure 4B**). However, *TrendCatcher* identified 978 DDEGs uniquely shown in the severe COVID-19 patient group. Importantly, these genes were strongly enriched for neutrophil-related biological pathways, including neutrophil activation and neutrophil mediated immunity. These severe-disease associated genes also included genes found in pathways such as myeloid cell differentiation, reactive oxygen species metabolic process, and positive regulation of cytokine production (**Figure 4B**).

**Figure 4.**
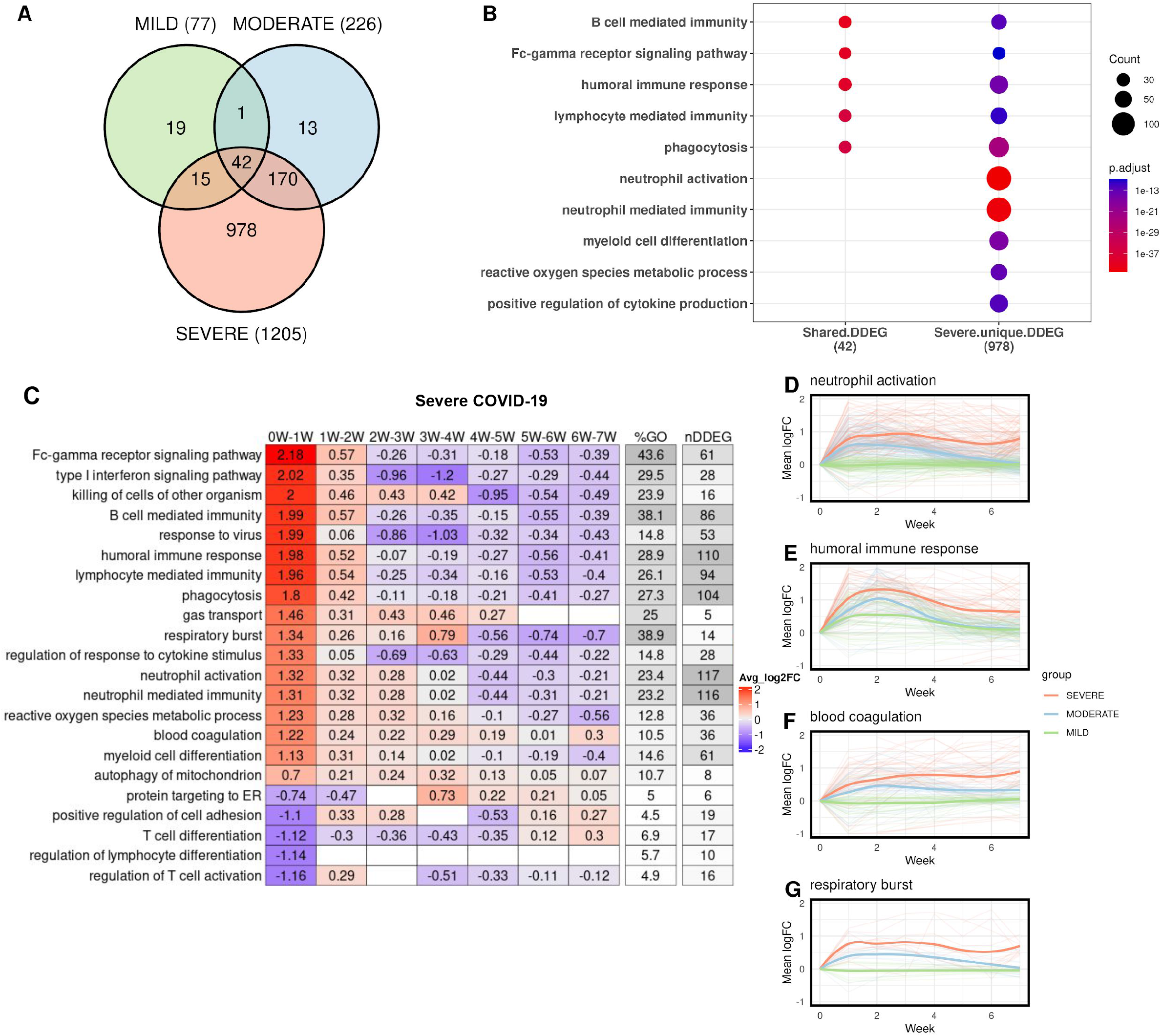
Temporal analysis of whole blood RNA-seq data in patients grouped according to disease severity. (A) Venn diagram of DDEGs identified from three COVID-19 severity groups. (B) Top GO enrichment from shared DDEGs across three groups compared with top GO enrichment from severe group. Dot size represents number of DDEGs enriched from the corresponding biological pathway. (C) *TimeHeatmap* of the top dynamic pathways from the severe group. Each column represents a time window, with W depicting week. (D-E) LOESS curve fitting of DDEGs identified in the severe COVID-19 group of the (D) neutrophil activation pathway, (E) humoral immune response pathway, (F) blood coagulation pathway and (G) respiratory burst pathway. Red curves represent the severe group, blue curves represent the moderate group, and green curves represent the mild group.

To systematically characterize which biological processes in whole blood RNA-seq samples were most dynamic in severe COVID-19 patients and how they progress over time, we applied the *TrendCatche*r’s *TimeHeatmap* function. As seen in the non-human primate PBMC and the human COVID-19 single cell RNA-seq datasets, we again observed the upregulation of genes involved in the response to virus, humoral immune response and type I interferon signaling pathway within Week 1 of disease onset (**Figure 4C**). Importantly, some dynamic biological responses were only enriched in the group of patients which subsequently progressed to severe COVID-19, including neutrophil activation (117 DDEGs, 23.4% of the corresponding GO term), blood coagulation (36 DDEGs, 10.5% of the corresponding GO term), and regulation of response to cytokine stimulus and respiratory burst (**Figure 4C, Supplemental Figure 6A,B**). For instance, neutrophil activation gene expression increased by a mean of 1.3 log2-units (approximately 2.5-fold increase in mean gene expression) within Week 1, increased continuously until Week 4, and only very gradually decreased in surviving patients by Week 7. However, the summation of the averaged log2 fold-change from the *TimeHeatmap* was larger than zero, which indicates that the neutrophil activation may not have returned fully to baseline levels by 7 weeks. To confirm this, we applied LOESS smooth curve fitting to all neutrophil activation DDEGs identified from the three severity groups. LOESS fitting confirmed that severe COVID-19 patients showed markedly higher neutrophil activation at the early stage of infection (Week 1- Week 2), and also remained highly activated even after 7 weeks. Such a persistent neutrophil activation was not observed in either mild or moderate COVID- 19 patient groups (**Figure 4D**).

We also observed severe COVID-19 patients showed evidence of greater humoral immune response gene upregulation (**Figure 4E**), as well as markedly higher upregulation blood coagulation genes (**Figure 4F**) and respiratory burst genes (**Figure 4G**), which remained upregulated even after Week 6. These were not found to be upregulated in patients with mild or moderate COVID-19 disease. These DDEGs and dynamic biological pathways may thus be suited to serve as early biomarkers that distinguish severe COVID-19 from mild and moderate COVID. All these dynamic gene signatures from neutrophil activation, blood coagulation and respiratory burst pathways were listed using traditional heatmap (**Supplemental Figure 7, Supplemental Figure 8A,B**).

### Early impaired IFN-I signaling in PBMC provides a hallmark of severe COVID-19

Next, to define the cell-type specific dynamic gene signatures and biological processes, we used *TrendCatcher* to analyze a human PBMC single cell RNA-seq dataset time course dataset in which patients were categorized as having either moderate or severe COVID-19 (15). Importantly, as this dataset only contained mRNA from peripheral blood mononuclear cells, it lacked mRNA from neutrophils. *TrendCatcher* generated “pseudo-bulk” mRNA profiles for each cell type in order to perform the analysis of gene expression dynamics in a cell-type specific manner. *TrendCatcher* identified more dynamic shifts in almost all cell types from severe COVID-19 patients than moderate groups (**Supplemental Table 1, 2**). To identify dynamic responsive processes unique to the severe group, we compared GO enrichment for each cell type between severe and moderate. As shown in **Figure 5A**, all innate immune cells and some adaptive immune cells (including B cells, CD8^+^ T cells) from moderate COVID-19 were highly enriched in type I interferon signaling, negative regulation of viral process and defense to virus, whereas this was not observed significantly enriched in severe COVID-19. Severe COVID-19 patients were highly dynamic in MAPK cascade, NF-κB signaling, T cell receptor signaling and positive regulation of cytokine production for both NK cells and CD4^+^ T cells. For monocytes and DC (dendritic cells), no uniquely enriched dynamic biological processes were observed in severe COVID-19 patients versus moderate COVID-19 patients.

**Figure 5.**
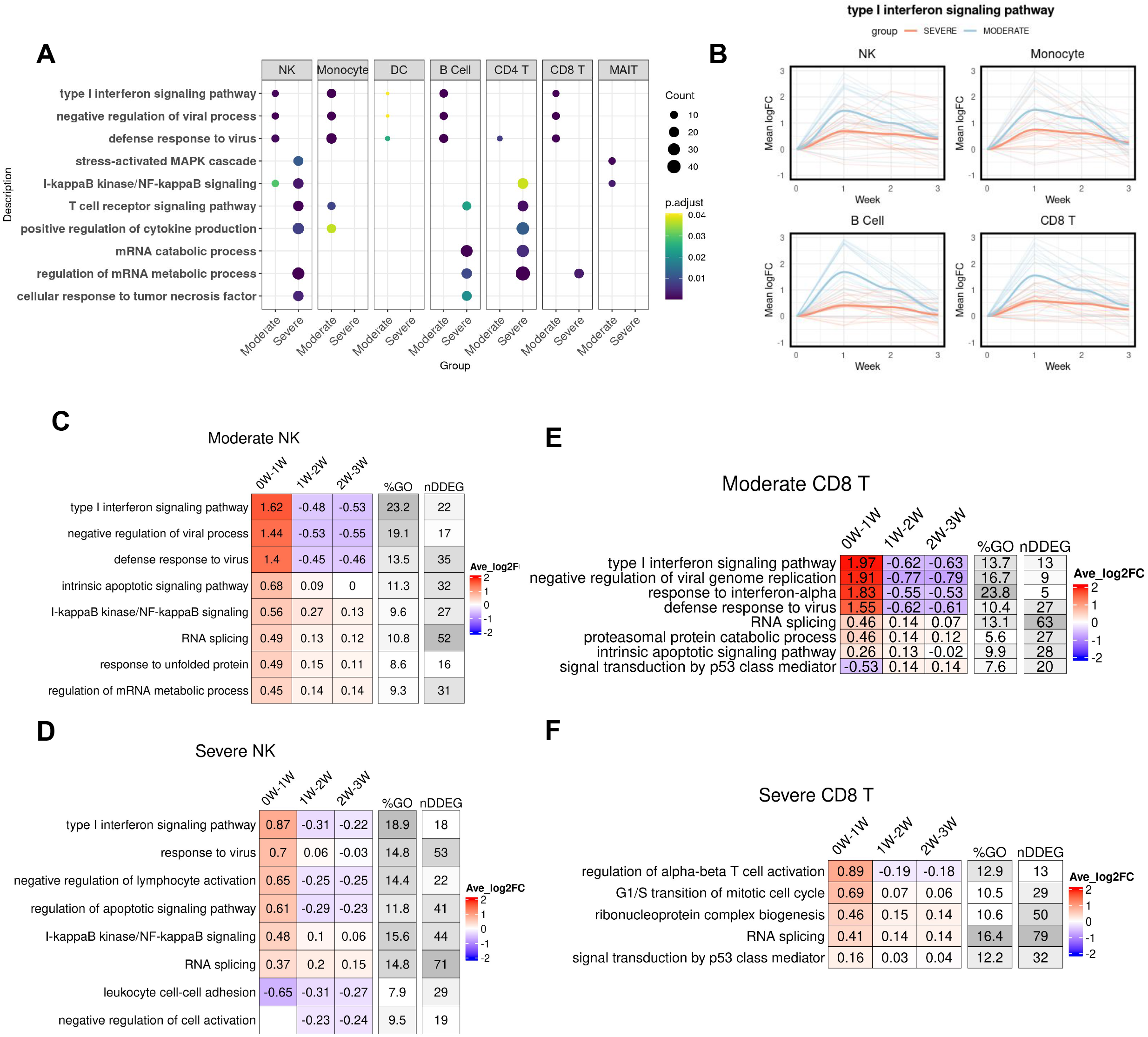
Temporal analysis of single cell RNA-seq data of PBMCs from patients with either moderate or severe COVID-19. (A) Dot plot showing GO enrichment comparison between severe COVID-19 and moderate COVID-19 for each cell type. The size of the dot represents gene count. (B) LOESS curve fitting on DDEGs identified from the type I interferon pathway using TrendCatcher from moderate COVID-19 and severe COVID-19. Blue color indicates moderate group and red color indicates severe group. (C) *TimeHeatma*p of NK cells from moderate COVID-19. (D) *TimeHeatmap* of NK cells from severe COVID-19. (E) *TimeHeatmap* of CD8^+^T cells from moderate COVID-19. (F) *TimeHeatmap* of CD8^+^T cells from severe COVID-19. Each column represents a time window with W depicting week.

Furthermore, we also found the extent of the interferon signaling response to be the key distinguishing feature between moderate and severe COVID-19. To quantitatively compare the trajectory differentiation of the type I interferon signaling pathway, we performed the LOESS smooth fitting to the DDEGs identified from this pathway. As shown in **Figure 5B**, there is a profound separation between these two groups, with early strong activation of IFN-I signaling in NK cells, monocytes, B cells and CD8^+^ T cells in moderate COVID-19 patients but not in severe COVID-19 patients. Patients with moderate COVID-19 showed activation of type I interferon signaling pathways whereas patients with severe COVID-19 had a blunted activation of type I interferon signaling. This is also shown in the cell-type specific *TimeHeatmap*. As shown in **Figure C,D**, although NK cells from both moderate and severe COVID-19 patients demonstrated activation of type I interferon response within the first week, moderate COVID-19 exhibited a stronger activation than severe group, with average 1.62 log2 fold-change compared to 0.87 log2 fold-change. In CD8^+^ T cells, only moderate COVID-19 groups were observed to have a strong type I interferon response within the first week. On the other hand, CD8^+^ T cells in patients who would go on to develop severe COVID-19, showed upregulation of cell proliferation and cell differentiation genes instead. Together, these data suggest that a robust early Type I interferon response in PBMCs is associated with reduced severity of COVID-19.

### *TrendCatcher* identifies metabolic gene expression shifts in NK cells as a hallmark response to COVID-19 vaccination

We next applied *TrendCatcher* to a longitudinal human PBMC single cell RNA-seq vaccination dataset (16), to provide insights into how the immune system physiologically responds to mRNA vaccines over time. The vaccination study collected single-cell PBMCs from 56 healthy volunteers vaccinated with the Pfizer-BioNtech at Day 0, 1, 2, 7, 21, 22, 28, and 42 after the vaccination. We processed each patient’s scRNA-seq individually. We clustered cells using the Seurat algorithm (17) and annotated the cell types using SingleR (18). Then we remove cell types containing less than 1,000 cells for each time point across all samples. As **Figure 6A** shown, it is the UMAP of one patient’s PBMCs single cell data at Day 0.

**Figure 6.**
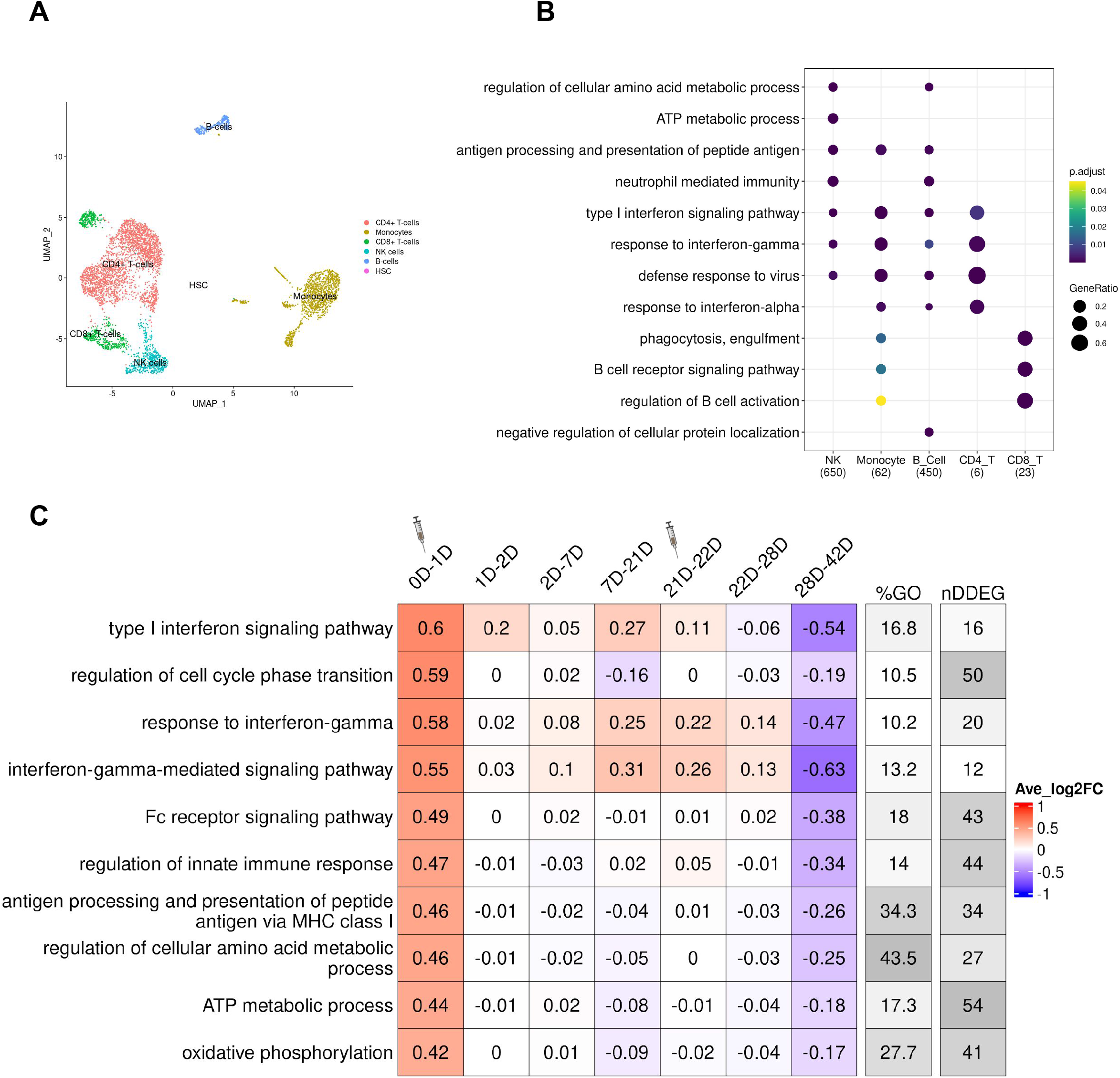
Temporal analysis of PBMC single cell RNA-seq data from human subjects receiving the SARS-CoV-2 mRNA vaccine. (A) UMAP of the single-cell transcriptional profile of patient 2047 at Day 0. Cell types were auto-annotated by SingleR. (B) Dot plot of comparison of the top GO terms enriched from cell-type specific DDEGs. *TrendCatcher* identified 650, 62, 450, 6 and 23 DDEGs from NK cells, monocytes, B cells, CD4^+^ T cells and CD8^+^ T cells. Size of the dot represents gene ratio from the enrichment analysis corresponding to each cell type. (C) T*imeHeatmap* of NK cells. Each column represents a time window measured by days. The second dose was administered at Day 21.

*TrendCatcher* identified 650 DDEGs in NK cells, 450 in B cells, 23 in CD8^+^ T cells, 62 in monocytes and only 6 in CD4^+^ T cells. This indicates NK cells exhibit a strong dynamic gene expression shift after vaccination. After comparing the GO enrichment analysis across cell types, we found NK cells gene expression shifts were enriched in metabolic processes in response to the Pfizer-BioNtech SARS-CoV-2 mRNA vaccine, such as regulation of cellular amino acid metabolism and ATP metabolism (**Figure 6B**). As shown in **Figure 6C**, a *TimeHeatmap* of NK cells shows 54 DDEGs, which account for nearly half of the ATP metabolic process pathway genes and exhibited a mean of 0.44 log2 fold upregulation after the first dose of vaccination. These findings indicate the reprogramming of metabolism in NK cells may be an indicator of an intact vaccine response. Another hallmark of the vaccine response was the activation of interferon pathways, including type I interferon signaling pathway and response to interferon-gamma (**Supplemental Figure 9A**). Importantly, the *TimeHeatmap* demonstrates that these gene expression shifts were very transient, usually decreasing by Day 2 or Day 7, and were again upregulated when the subjects received the booster vaccine at Day 21. Interferon pathways showed an average increase of 0.71 and 1.42 log2 fold-change. In adaptive immune cells, we also observed strong type I interferon signaling pathways activated in both B cells and CD8^+^ T cells (**Supplemental Figure 9B,C**). Additionally, B cells demonstrated upregulation of antigen processing and protein synthesis (**Supplemental Figure 9B**), which were upregulated only for the first day, and then began decreasing. It is noteworthy that early upregulation of interferon pathways in the PBMCs of vaccine recipients mirrors that seen in patients with moderate COVID-19 infection, consistent with the notion that early and transient interferon upregulation is a hallmark of a healthy immune response to the SARS-CoV-2 infection.

## Discussion

Temporal analysis of gene expression is gaining importance in the analysis of complex dynamic processes such as disease progression. Besides gene dynamic pattern characterization, time-course gene expression data are also used to infer regulatory and signaling relationships among genes (19, 20). Integrating with other different types of measurements, such as pathology and infection over time helps disentangle the complex dynamic processes and possible underlying mediators (21). Thus, accurate identification of dynamic differentially expressed genes (DDEGs) in a time course RNA-seq or single cell RNA-seq study can help identify time-dependent disease mechanisms, adaptive and maladaptive molecular signatures as well as potential biomarkers that may be associated with disease severity.

In recent years, tools have been implemented to characterize time course RNA-seq data; however, these tools were focused on either bulk RNA-seq datasets (3, 6, 7) and few infer trajectories from scRNA-seq (22, 23). Compared to methods that analyze time course data without considering the sequential nature of time points, modeling time as a continuous function avoids a relative loss of statistical testing power, especially when many multiple time points were studied. There is also a need for methods which combine DDEGs identification with the visualization of dynamic pathway shifts.

In this study, we developed an open-source R-software package to perform temporal analysis in longitudinal bulk RNA-seq and scRNA-seq datasets. *TrendCatcher* can identify DDEGs and infer the trajectories for each DDEG. By combining a time-window sliding strategy on inferred gene trajectories and leveraging annotated biological pathway databases, *TrendCatcher* can infer and visualize the most dynamic biological pathways in response to the external stimuli. Furthermore, by utilizing the *TimeHeatmap* function, *TrendCatcher* can help researchers identify the magnitude and dynamic nature of pathways shifts. Using simulated datasets for benchmarking, *TrendCatcher* achieved higher accuracy (AUC = 0.9) compared to other commonly used methods for the analysis of temporal gene expression datasets, when analyzing three or more time points. The advantage of using *TrendCatcher* was especially apparent in the setting of biphasic or multiphasic temporal trajectories, in which gene expression levels can change their trend of upregulation or downregulation during the time course.

Despite the extraordinary success of rapidly developed and deployed mRNA vaccines against SARS-CoV-2, the ongoing COVID-19 pandemic remains a major global health problem in part due to the emergence of newer highly contagious SARS-CoV-2 variants of concern as well as vaccine hesitancy. This requires the identification of novel mechanistic targets, especially in vulnerable patients who have a high risk of developing severe COVID-19. One of the key pathogenic factors driving COVID-19 severity is the profound immune dysregulation observed in patients with severe COVID-19 that can result in respiratory failure (24–27). Human immune responses to SARS-CoV-2 infection are highly dynamic and time dependent, requiring upregulation as well as downregulation of distinct immune signaling pathways at the appropriate times to ensure optimal host defense (24–27). Understanding the dynamics of the COVID-19 immune response could form the basis of developing therapies that are appropriate for a given time window (28). Personalized medicine or precision medicine are gaining traction by developing therapies tailored to patients based on their genotype and phenotype, but personalization likely also requires tailoring therapies based on the temporal phase of a disease. These studies highlight the need for time-course or longitudinal analyses of the host responses to the SARS-CoV-2 infection.

In this study, we applied *TrendCatcher* to systematically analyze sequential blood samples from either non-human primates infected with SARS-CoV-2, human patients with COVID-19 of varying severity or human subjects who received a SARS-CoV-2 mRNA vaccine. *TrendCatcher* identified dynamic gene expression and biological pathway signatures for (a) SARS-CoV-2 infection progression over time, (b) severe COVID-19 vs. moderate or mild COVID-19 and (c) COVID-19 vaccine recipients versus control. *TrendCatcher* established response to virus, humoral immune response and type I interferon signaling pathway activation across peripheral blood cell types in mild, moderate and severe COVID-19. However, we found temporal patterns of gene expression shifts that were unique in the severe COVID-19 patients. Severe COVID-19 was associated with marked activation of neutrophils and upregulation of coagulation pathways as well as blunted type I interferon signaling as early as week 1 in the peripheral blood of patients. Importantly, severe COVID-19 was associated with persistent activation of neutrophils and genes regulating coagulation for as long as 6 weeks, underscoring that the importance of the temporal analysis by *TrendCatcher* which identified hallmarks of severe COVID-19 in peripheral blood samples.

Our findings complement recent studies which implicate neutrophils in the excessive inflammation, coagulopathy, immunothrombosis and organ damage which characterize severe COVID-19 (29–33). Neutrophils are particularly active in highly vascularized organs, such as lungs and kidneys, which are prime targets of SARS-CoV-2 induced injury in COVID- 19 (34). Dysregulated neutrophil responses, such as prolonged activation may cause damage to vessels and underlying parenchymal (35–37). Studies also show that activated neutrophils through Toll-like receptors, chemokine- and cytokine receptors can stimulate the neutrophil extracellular trap (NET) formation (29). And recent studies showed the disbalance between NET formation and degradation can trigger immunothrombosis and tissue injury (38, 39). Excessive activation of macrophages and adaptive immune cells in severe COVID-19 which can form vicious cycles of positive feedback circuits has also been demonstrated (40, 41), but less is known about how early neutrophils are activated in disease progression because many of the studies on neutrophils in clinical COVID-19 or animal models of severe SARS-CoV-2 focused on the late stages of the disease as well as the neutrophils in the lung tissue. In our study, we used a systematic temporal analysis and observed that profound neutrophil activation in the peripheral blood, which was predominant in the severe COVID-19 group, occurs as early as week one after diagnosis or symptoms persists even after six or seven weeks in surviving patients.

Our observation of early upregulation of coagulation genes in the whole blood transcriptomes of severe COVID-19 patients also points to another feature that is associated with severe COVID-19, thrombotic and embolic complications such as strokes (13, 42, 43). Recent studies have found the procoagulant changes in endothelial cells underly the coagulopathy in severe COVID-19 (44). Endothelial dysfunction in COVID-19 patients may be exacerbated through inflammatory cytokines and neutrophil extracellular traps, thus pointing to interactions between circulating or recently recruited neutrophils and the vessel wall endothelial cells that are in contact with circulating immune cells as drivers of such coagulopathic manifestations (45–47). We believe the analysis of the RNA-seq and scRNA-seq data support neutrophil activation and upregulation of coagulation genes as defining features of severe COVID-19, highlighting their role as early biomarkers to provide prognostic information and thus optimally treat patients who have a higher likelihood of progressing to severe disease. Importantly, our results also raise questions about the role of neutrophils and of coagulation in what is referred to as “long COVID” or “post-acute sequelae of COVID” (48–50), because we observed that the activation persists for as long as six to seven weeks after the initial infection.

Our temporal analysis showed that severe COVID-19 single-cell PBMCs were characterized by impaired type I interferon signaling at the onset of infection (Week 1), compared to the moderate COVID-19 group. This impaired type I interferon signaling was identified in innate cells, such as NK cells and monocytes, and adaptive cells, such as B cell and CD8^+^ T cells. Type I interferons, which are essential for antiviral immunity (51, 52), have been shown to be upregulated in COVID-19 (53, 54). Other studies also suggested that impaired type I interferon signaling may promote severe COVID-19 and that interferon therapy could be used as therapy in severe COVID-19 (55). However, since these studies were not designed for longitudinal analyses, the timing of when to intervene on IFN signaling was not clear. Traditional analyses of gene expression data are not suited for the identification of temporal windows and thus do not address whether impaired IFN production is present at the onset of infection, whether it was delayed, or whether IFN production is exhausted after an initial activation (54). *TrendCatcher* addressed this question by showing the dynamic timing of type I IFN from both the moderate group and severe group.

In our COVID-19 vaccination single cell PBMC temporal data analysis, we also identified prominent metabolic shifts in NK cells. Cellular metabolism is recognized as an important factor that can determine the fate and function of lymphocytes (56, 57). Certain metabolic pathways have been shown to have direct immunoregulatory roles (56). Activated NK cells engage in a robust metabolic response, which is required for normal effector functions (58), and our data suggests that assessing NK metabolic shifts may be an intriguing alternative to assessing vaccine responsiveness, which does not rely on antibody levels which can be highly variable in laboratory assays.

In conclusion, we have developed the novel *TrendCatcher* R package platform designed for time course RNA-seq data analysis which identifies and visualizes distinct dynamic transcriptional programs. When applied to real whole blood bulk RNA-seq time course datasets from COVID-19 patients, we observed that patients with severe COVID-19 showed gene expression profiles consistent with profound neutrophil activation and coagulopathy early during the progression of the disease (starting from the first week of symptom onset). This indicates that neutrophil activation may be a key early biomarker and possible mediator of severe COVID-19. The distinct kinetics of gene expression shifts identified by *TrendCatcher* in COVID-19 patients may allow for the identification of therapeutic targets during distinct temporal phases.

## Author contributions

X.W. and J.R. designed the study, X.W. performed the analyses and wrote the first draft of the manuscript, J.R. co-wrote the initial draft of the manuscript. M.S. and Y.D. provided critical revisions of the manuscript.

## Acknowledgments

The studies were supported by NIH grants R01-HL149300 (to JR), P01-HL151327 (to JR), and T32-HL139439 (to MS).

## Materials and Methods

### TrendCatcher framework

The main components of the *TrendCatcher* framework are shown in **Figure 1**. *TrendCatcher* requires two main inputs: the raw count table C of a temporal study with a dimension of *m* × *n*, where *m* denotes the number of genes and *n* denotes the number of samples, and a user-defined baseline time variable *T*, such as “0 hour”. Since samples may have different sequencing depths and batch effect, *TrendCatcher* integrates with limma (5) and provides preprocessing steps, such as batch correction and normalization. For single-cell RNA sequencing datasets, *TrendCatcher* extracts cells for each cell type annotated in the meta data slot of Seurat object and convert it into cell-type specific “pseudobulk” time course RNA library. Based on a user-specified threshold, relatively low abundant genes are removed from the count table, and reads are normalized and batch effects are removed. *TrendCatcher*’s core algorithm is composed of five main steps: (a) baseline fluctuation confidence interval estimation, (b) model dynamic longitudinal count, (c) time point dynamic p-value calculation, (d) gene-wise dynamic p-value calculation, and (e) break point screening and gene-wise dynamic pattern assignment. Mathematical details will be expanded in the following sections. For the output of *TrendCatcher*, there are mainly two components: a master table and a set of functions for versatile visualization purposes. The master table contains all the dynamic details of each single gene, including its dynamic p-value, its break point location time, and its dynamic trajectory pattern. In addition to the master table, *TrendCatcher* produces five main types of visualizations: (a) a figure showing the observed counts and fitted splines of each gene, (b) genes trajectories from each trajectory pattern group (c) a hierarchical pie chart that represents trajectory pattern composition, (d) a *TimeHeatmap* to infer trajectory dynamics of top dynamic biological pathways and (e) a two-sided bar plot to show the top most positively and negatively changed (averaged accumulative log2FC) biological pathways.

### Baseline fluctuation confidence interval estimation

We assumed that the observed number of RNA-seq reads count from the baseline time (e.g., t=0 hour) *X_i,t_baseline__* was generated from a negative binomial distribution (Equation 1):

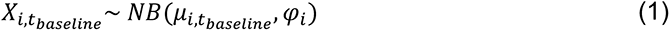

Where gene *i* = 1, …, *n* and *μ_i,t_baseline__* is the mean count of gene *i* at baseline time, and *φ_i_* is the dispersion factor. First, the dispersion factor *φ_i_* was pre-estimated as a constant hyperparameter for each gene with DESeq2 (3), as shown in Equation 2. Here, *σ_i_*(*t*) is the variance at time t.

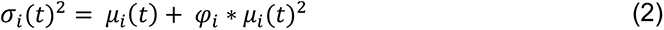

Then, *μ_i,t_baseline__* was estimated using maximum likelihood from Equation 3.

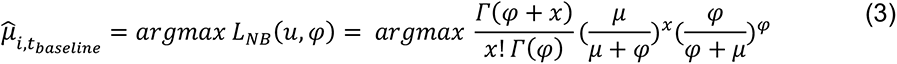

Based on the negative binomial distribution and the estimated mean count for baseline time, we constructed the 90% confidence interval [*X_i,t_baseline__*_0.05_, *X_i,t_baseline__*_0.95_] as the baseline count fluctuation interval (Equation 4(a) and Equation 4(b)).

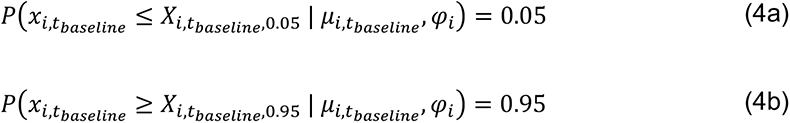

### Model dynamic longitudinal count estimation

To model the time-dependent gene expression value, we applied a smoothing spline ANOVA model (59, 60) with a negative binomial family constraint to fit the reads from samples across non-baseline multiple time points. The random variable *X_i,t(t_*_≠*tbaseline)*_ is assumed to follow the NB distribution in Equation 5, with a positive integer *α* represents number of failures before the *α*th success in a sequential of Bernoulli trials, and *p*(*t*) ∈ (0, 1) represents the success probability.

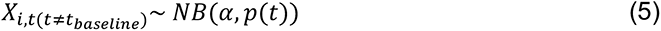

The log-likelihood given a time-course observed count *x* = [*x_i,t_*_(*t*≠*t_baseline_*)_}*_i_*_=1,…,*n*; *t*=1,…,*T*_ is calculated as Equation 6.

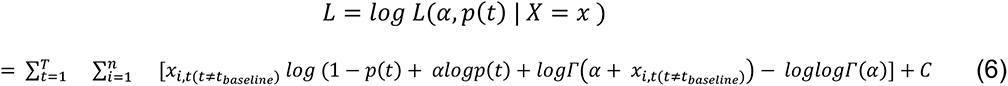

Taking the logit link and model time effect, we define the logit link *η*

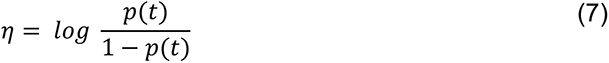

To allow for flexibility in the estimation of *η*, and find the best trade-off between goodness of fit and the smoothness of the spline curve, soft constraints of the form *J*(*η*) is added to the minus log-likelihood, with the smoothing parameter *λ* > 0.

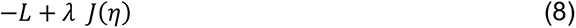

The solution to the optimization of Equation 8 leads to a smoothing fitting to the reads from samples across different non-baseline time points. The estimated mean of *μ_i,t_*_(*t*≠*tbaseline*)_ can be estimated using Equation 9.

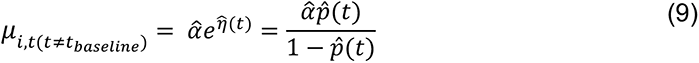

### Gene’s dynamic p-value calculation

To calculate gene’s non-baseline dynamic signal significance, each gene’s non-baseline estimated mean count *μ_i,t_*_(*t*≠*t_baseline_*)_ was tested against the baseline fluctuation interval.

Based on Equation 10(a) and Equation 10(b), for each gene at each single non-baseline time point, a dynamic time p-value was calculated.

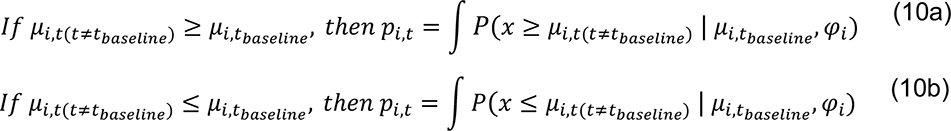

Then, we applied Fisher’s combined probability test method to calculate a gene-wise dynamic p-value (Equation 11):

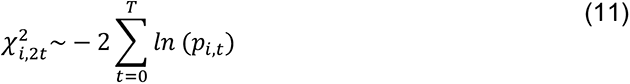

### Trajectory pattern assignment

First, we connect all the significant dynamic signal time points, with a p-value threshold less than 0.05. Then we applied a break point searching strategy to capture the gene expression change trend. The definition of break point is defined using Equation 12(a) and Equation 12(b).

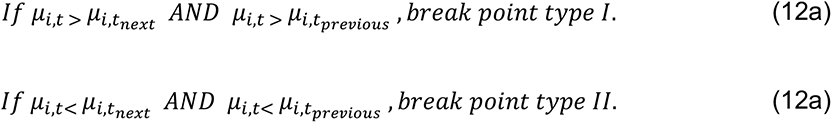

There are two types of break points: type I means gene up-regulated followed by a down regulation, and type II means gene expression level down regulated and then followed by an upregulation. By screening along the break point, the master-pattern and sub-pattern were assigned to each gene.

### *TimeHeatmap* enrichment analysis and Two-sided bar plot

To build the *TimeHeatmap* for visualizing the biological pathway enrichment change over time, we designed a window-sliding strategy to capture all the up-regulated or down-regulated genes within each time interval. If we denote time vector as *t_i_*; and *j* ∈ 1, …, *T*, each time interval is denoted as [*t_j_*_-1_, *t_j_*]. We found *N_up_* regulated genes and *N_down_* down-regulated genes within the time window [*t_j_*_-1_, *t_j_*], then Fisher’s exact test was performed to obtain the Gene Ontology (GO) term enrichment with the corresponding time interval for up-regulated genes and down-regulated genes separately. Users can select the top most enriched biological pathways (ranked by enrichment p-value) for each time interval (by default is 10). Then for each selected GO term within the corresponding time window, we calculated the averaged log 2 fold-change (Avg_log2FC) of all the DDEGs from this GO term. A series of Avg_log2FC values over time characterize the trajectory dynamics of the corresponding biological pathway, it is defined as biological pathway trajectory inference in this study. The summation of the series of Avg_log2FC estimates the averaged accumulative log 2 fold-change (GO_mean_logFC) for the corresponding GO term. *TrendCatcher* ranks biological pathways based on their dynamic magnitude inferred from the GO_mean_logFC value. Users are free to choose the top most positively and negatively changed (averaged accumulative log2FC) biological pathways. Besides GO enrichment analysis (61), *TrendCatcher* also packages Enrichr (62) biological pathway databases. Visualization is constructed using the ComplexHeatmap (63) package.

### Simulated dataset

To mimic the real biological RNA-seq dataset, we only allowed ∼10% of the genes to be dynamic responsive genes. In this study, we embedded 5 different types of trajectories into the temporal RNA-seq simulated datasets, including non-dynamic trajectory (∼90%), monotonic trajectory (∼2.5%), bi-phase trajectory (∼2.5%), and multimodal trajectory (∼5%) including 2 break-point and 3 break-point trajectory. Each type of trajectory was constructed by adding negative binomial distribution noise to the embedded trajectory count. For example, for monotonic trajectory, we defined the first and last time-points’ RNA expression level change to be 0.5-2 log2 fold-change, then we added negative binomial distribution noise to the embedded monotonic trajectory. To check how the AUC changes as time-course study extends longer, we sampled 3, 5, 7, 9 and 11 different number of time points with evenly time intervals, and randomly sampled 3 replicates for each time point. To validate how the prediction AUC varies across different types of trajectories, we embedded ∼10% of the genes to have non-dynamic trajectories, and ∼90% of the genes have a specific dynamic trajectory (monotonic, bi-phase or multimodal).

### Pseudobulk RNA library construction

To construct “pseudo-bulk” RNA library from single cell RNA-seq datasets, all cells for each cell type in a given sample were computationally “pooled” by summing all counts for a given gene. Since pseudobulk libraries composed of few cells are not likely modeled properly, we removed cell types containing less than 1k cells in this study. Lowly expressed genes were removed for each cell type as well, using the filterByExpr function from *edgeR* R package (4). Gene counts were transformed using the log of the counts per million (cpm) and library size was normalized using calcNormFactors function with the method = “RLE”.

### Gene set enrichment of DDEGs

Gene sets enrichment analysis in this study was performed using clusterProfiler (61) R package, and enrichment comparison from multiple groups were visualized using the compareCluster function.

---

**Table 1.** Datasets presented in this study

| Dataset | Organism | Number of time points | Study | Figure | Dataset Source | Reference |
| --- | --- | --- | --- | --- | --- | --- |
| Bulk Peripheral Blood<br>(Aid et al.) | Macaca mulatta | 7 | Covid vs. Normal | Figure 2 | GSE156701 | (13) |
| scRNA PBMC<br>(Zhu et al.) | Homo Sapien | 5 | Covid vs. Normal | Figure 3 | https://db.cngb.org/search/project/CNP0001102/ | (14) |
| Bulk Peripheral Blood<br>(Bergamaschi et al.) | Homo Sapien | 8 | Severe vs. Moderate | Figure 4 | https://www.covid19cellatlas.org/patient/citiid/ | (11) |
| scRNA PBMC<br>(Liu et al.) | Homo Sapien | 4 | Severe vs. Moderate | Figure 5 | GSE161918 | (15) |
| scRNA PBMC<br>(Arunachalam et al.) | Homo Sapien | 8 | Vaccination | Figure 6 | GSE171964 and GSE169159 | (16) |

**Supplement Table 1.** Number of DDEGs for each innate immune cell type identified by *TrendCatcher*. Using FDR less than 0.05.

| Innate immune cells |  |  |  |  |  |
| --- | --- | --- | --- | --- | --- |
| NK |  | Monocyte |  | DC |  |
| Moderate | Severe | Moderate | Severe | Moderate | Severe |
| 708 | 1,179 | 1,479 | 1,845 | 22 | 17 |

**Supplement Table 2.** Number of DDEGs for each adaptive immune cell type identified by *TrendCatcher*. Using FDR less than 0.05.

| Adaptive immune cells |  |  |  |  |  |  |  |
| --- | --- | --- | --- | --- | --- | --- | --- |
| B |  | CD4<sup>+</sup> T |  | CD8<sup>+</sup> T |  | MAIT |  |
| Moderate | Severe | Moderate | Severe | Moderate | Severe | Moderate | Severe |
| 969 | 1,344 | 190 | 1,060 | 634 | 817 | 71 | 13 |

**Supplementary Figure 1, related to Figure 1.**
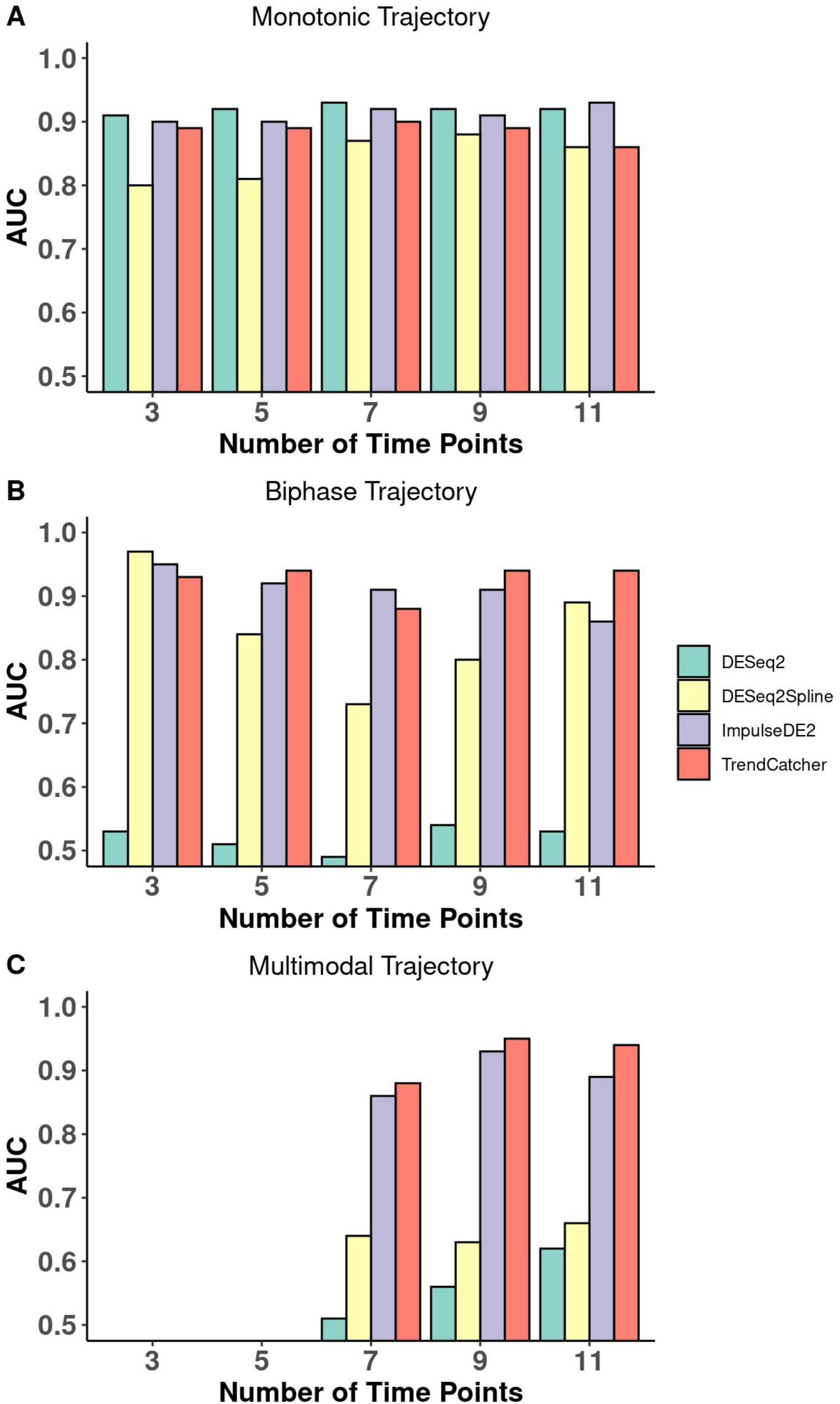
TrendCatcher prediction performance on different types of trajectories. (A) Prediction performance of *TrendCatcher* across varying numbers of time points for monotonic trajectories. (B) Prediction performance of *TrendCatcher* across varying numbers of time points for biphasic trajectories. (C) Prediction performance of *TrendCatcher* across varying numbers of time points for multimodal trajectories. DESeq2 is shown in green, DESeq2Spline is shown in yellow, ImpulseDE2 is shown in purple, *TrendCatcher i*s shown in red.

**Supplementary Figure 2, related to Figure 2.**
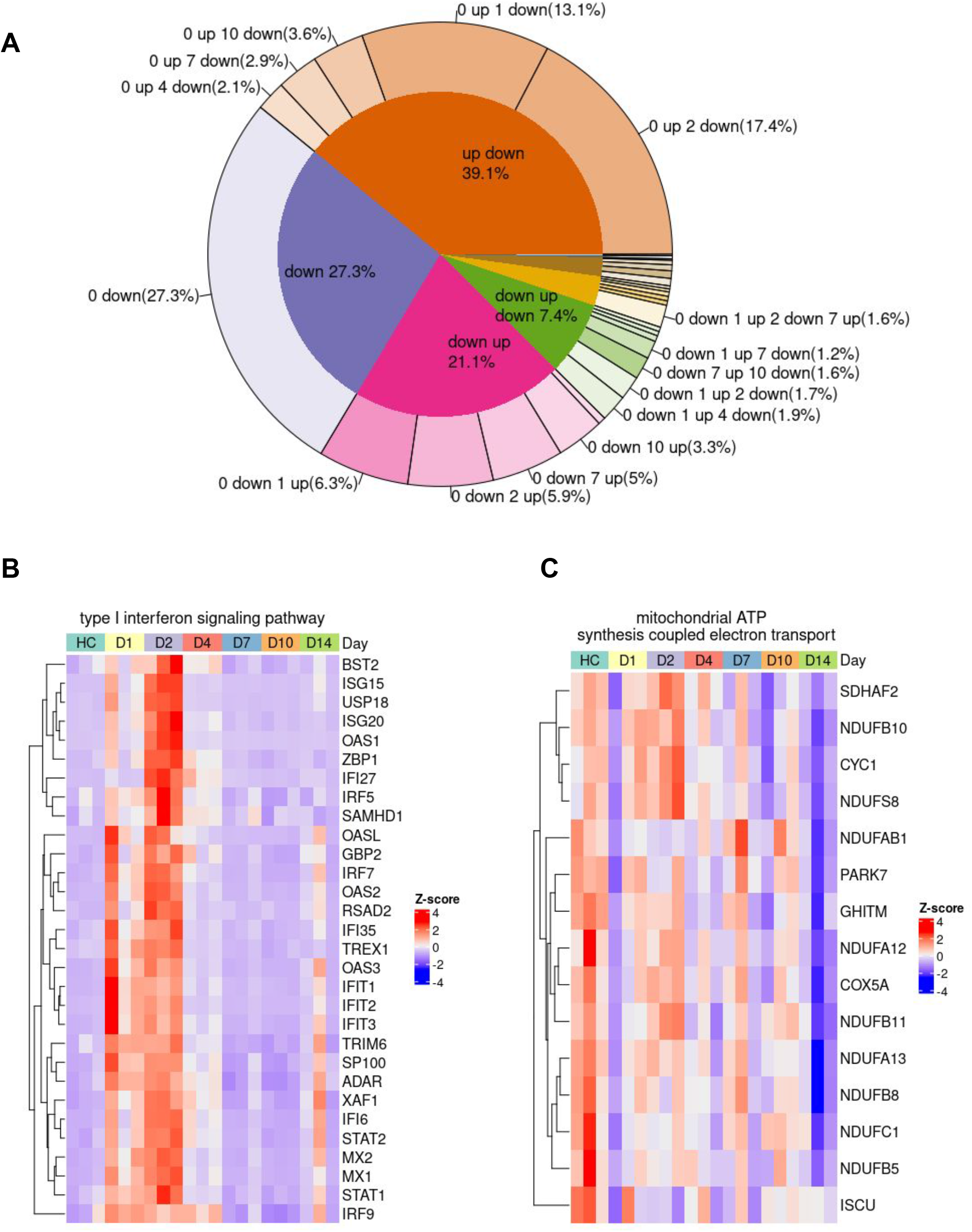
Trajectory pattern composition of DDEGS and dynamic gene signatures from highlighted pathways. (A) Hierarchical pie chart shows the composition of trajectory patterns of DDEGs identified in a non-human primate bulk peripheral blood mRNA (Aid et al.). (B) Dynamic gene signatures for SARS-CoV-2 infection from type I interferon signaling pathway. (C) Dynamic gene signatures response for SARS-CoV-2 infection from mitochondrial ATP synthesis coupled electron transport pathway. Color represents normalized z-score for gene expression.

**Supplementary Figure 3, related to Figure 3.**
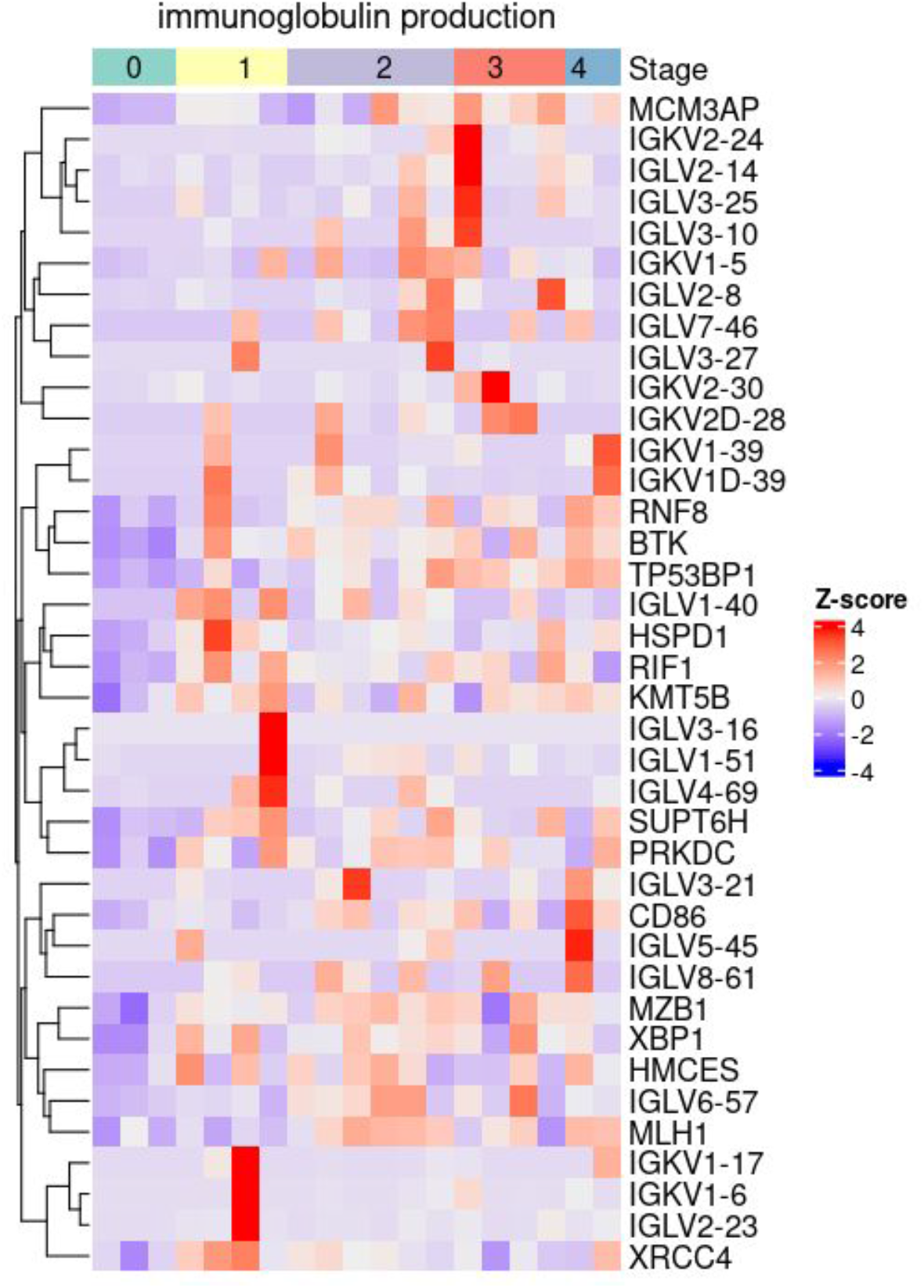
Dynamic gene signatures of immunoglobulin pathway. Heatmap showing B cell’s 38 DDEGs identified from immunoglobulin production process. Color represents normalized z-score of gene expression.

**Supplementary Figure 4, related to Figure 3.**
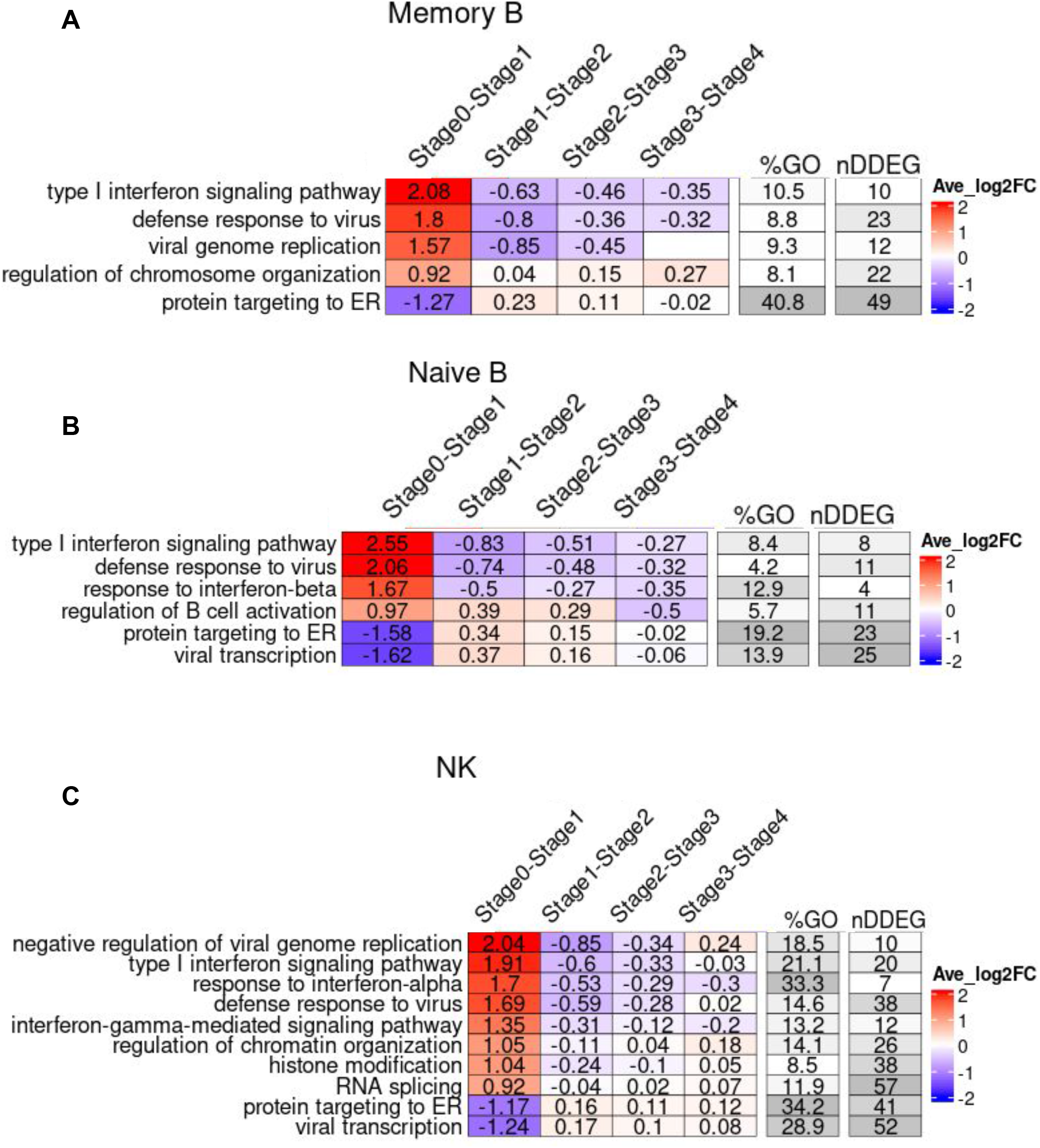
TimeHeatmap of B cells and NK cells. **(A)** *TimeHeatmap* of Memory B cells. (B) *TimeHeatmap* of Naive B cells. (C) *TimeHeatmap* of NK cells.

**Supplementary Figure 5, related to Figure 3.**
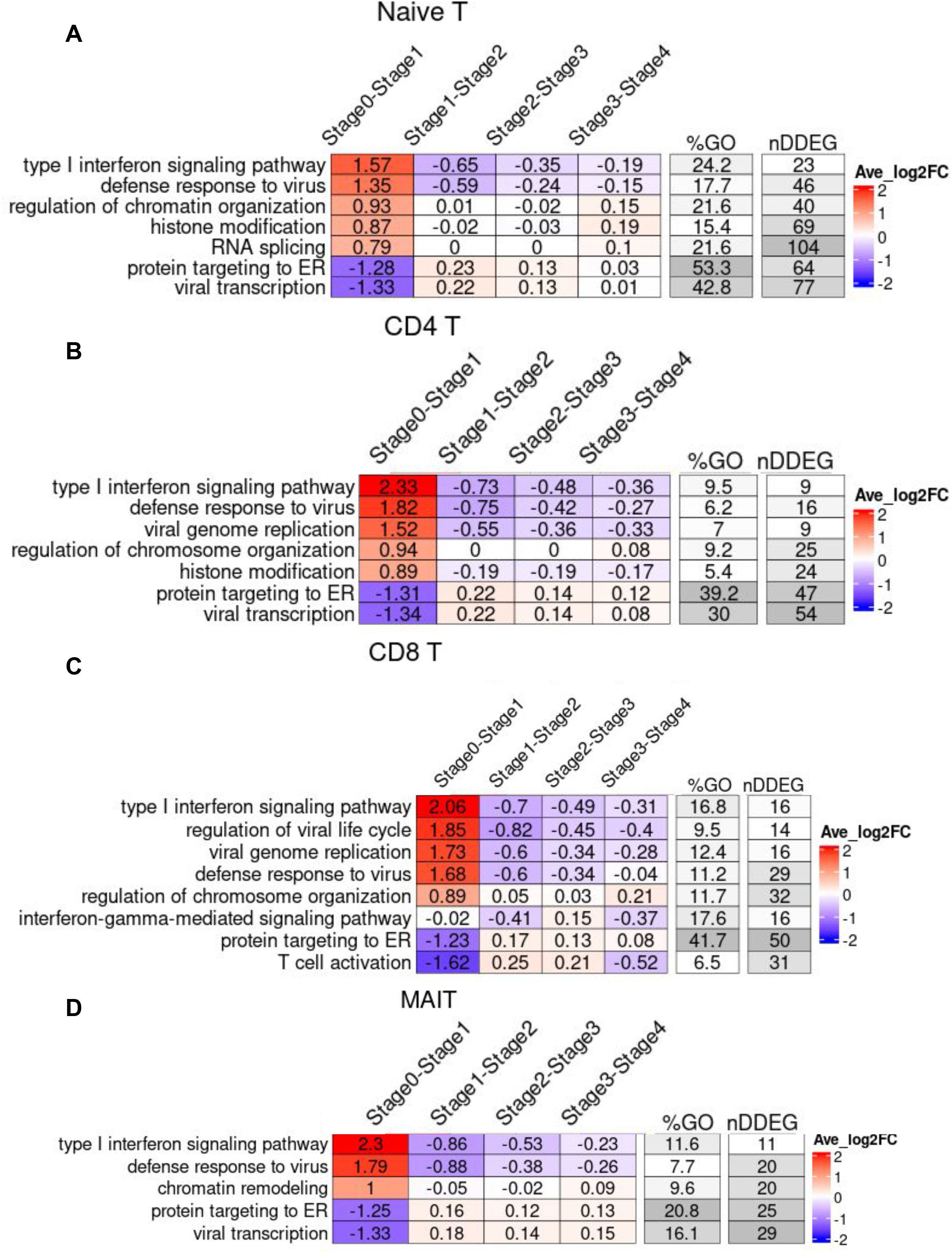
TimeHeatmap of T cells. (A) *TimeHeatmap* of Naive T cells. (B) *TimeHeatmap* of CD4^+^ T cells. (C) *TimeHeatmap* of CD8^+^ T cells. (D) *TimeHeatmap* of MAIT cells.

**Supplementary Figure 6, related to Figure 4.**
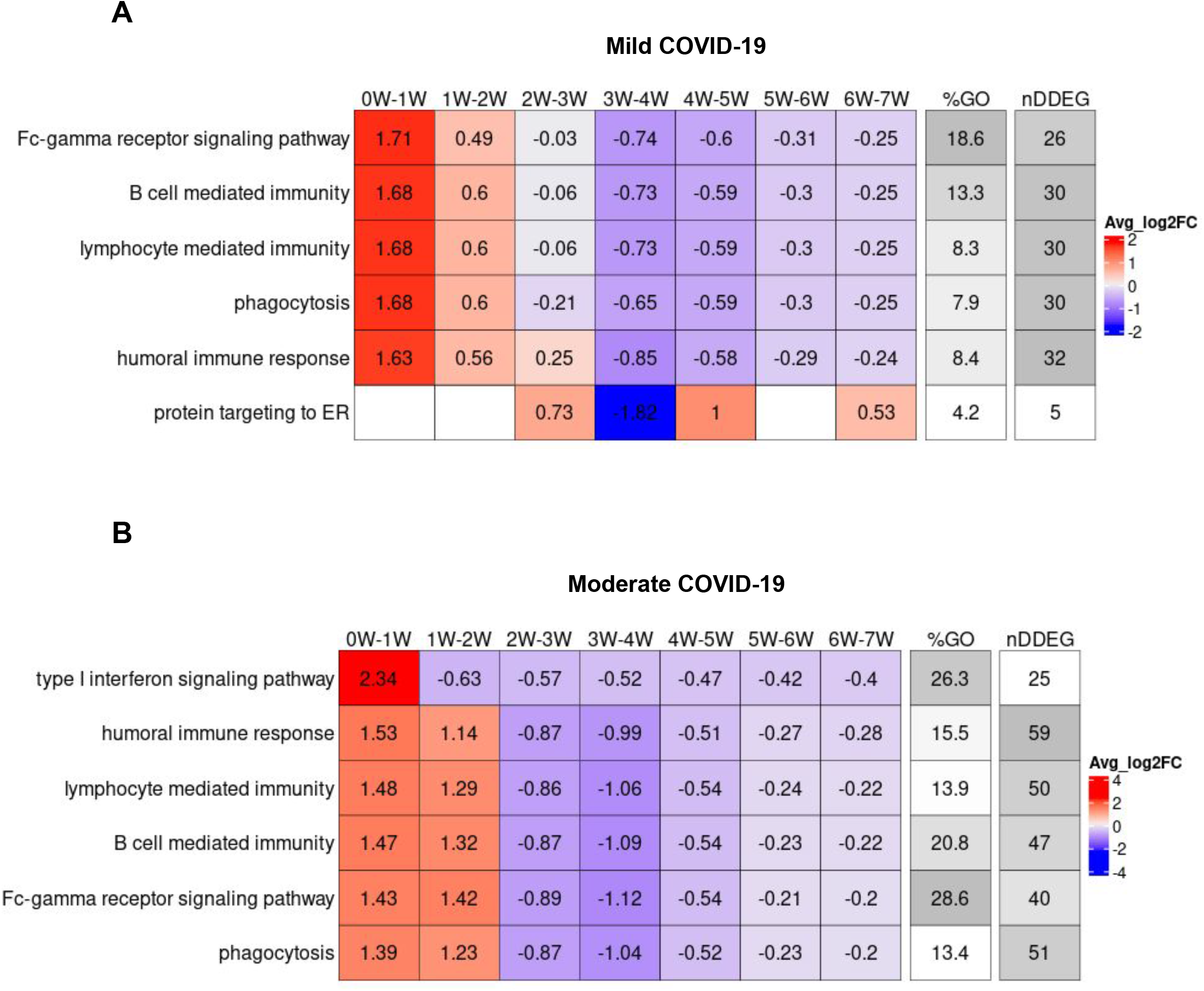
TimeHeatmap of mild and moderate COVID-19. (A) *TimeHeatmap* of top dynamic GO terms found in severe COVID-19 shown in mild group. (B) *TimeHeatmap* of top dynamic GO terms found in severe COVID-19 shown in moderate group.

**Supplementary Figure 7, related to Figure 4.**
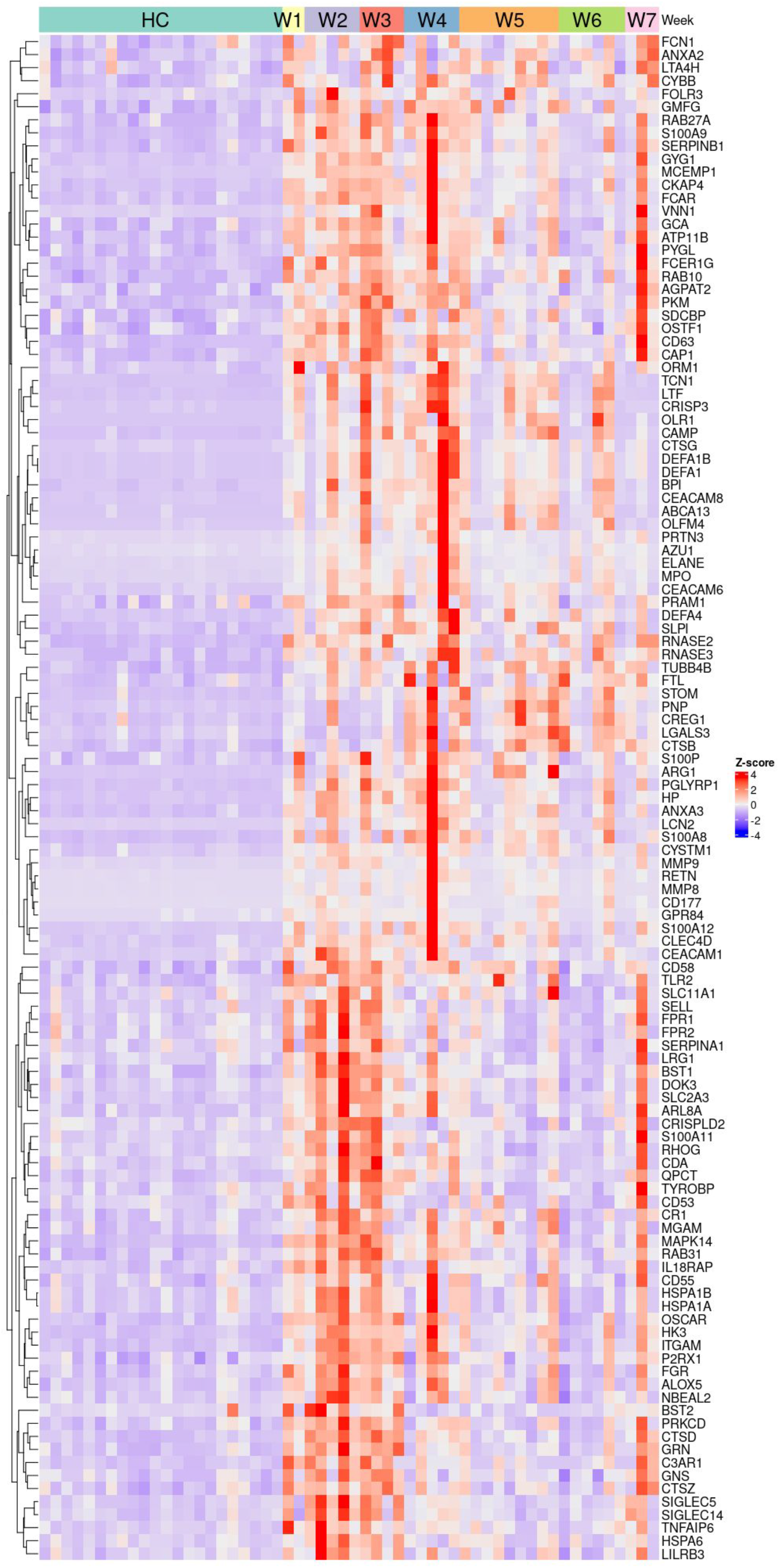
Dynamic gene signatures of neutrophil activation from severe COVID-19. Heatmap of severe COVID-19 DDEGs identified from the neutrophil activation pathway. The heatmap is ordered by week following infection with healthy controls (HC) on the left. Color represents Z-score normalized expression values. Each column represents one sample.

**Supplementary Figure 8, related to Figure 4.**
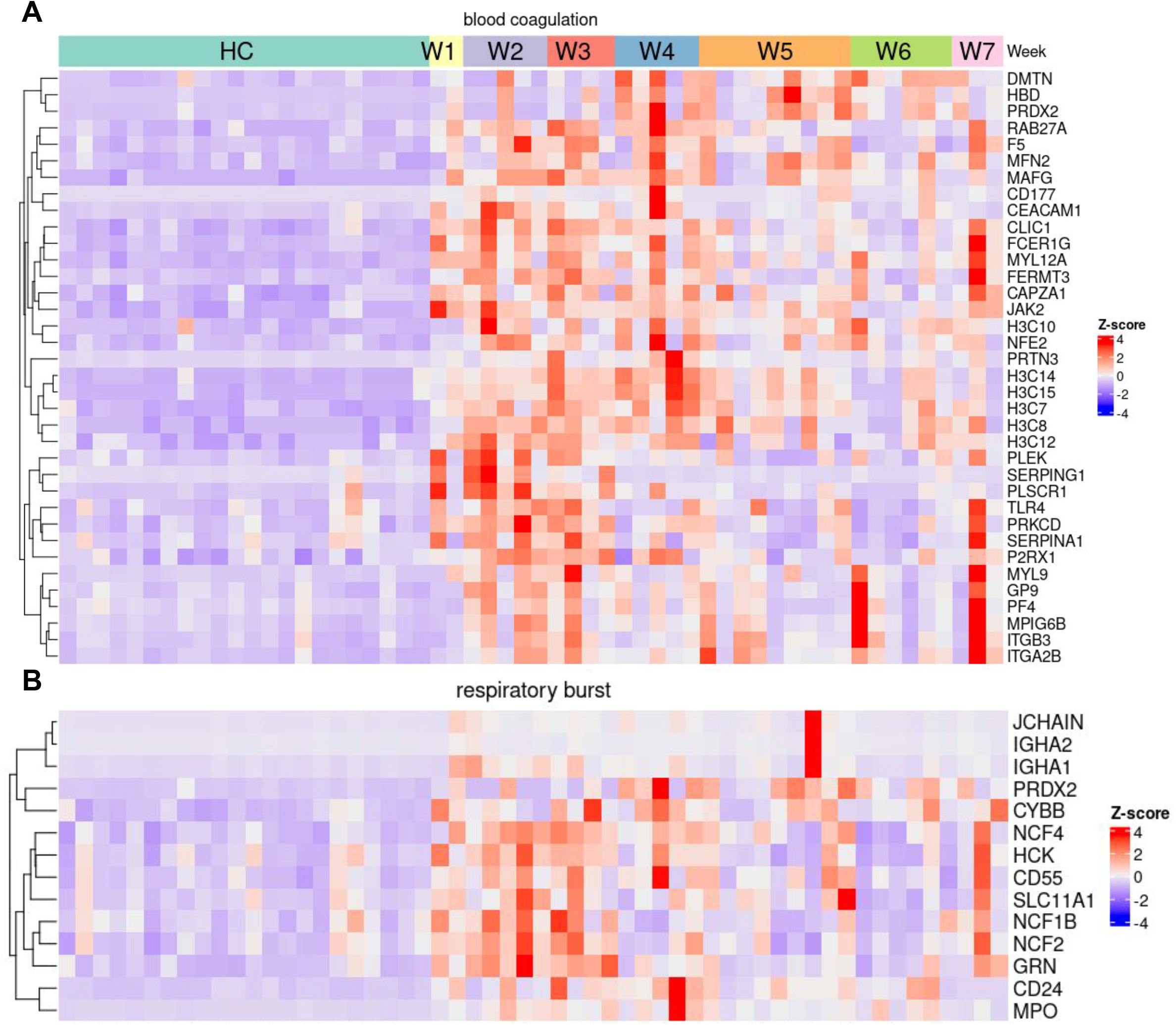
Dynamic gene signatures of blood coagulation and respiratory burst pathways from severe COVID-19. (A) Severe COVID-19 DDEGs identified from the blood coagulation pathway. (B) Severe COVID-19 DDEGs identified from respiratory burst pathway. The heatmap is ordered by week following infection with healthy controls (HC) on the left. Color represents Z-score normalized expression values.

**Supplementary Figure 9, related to Figure 6.**
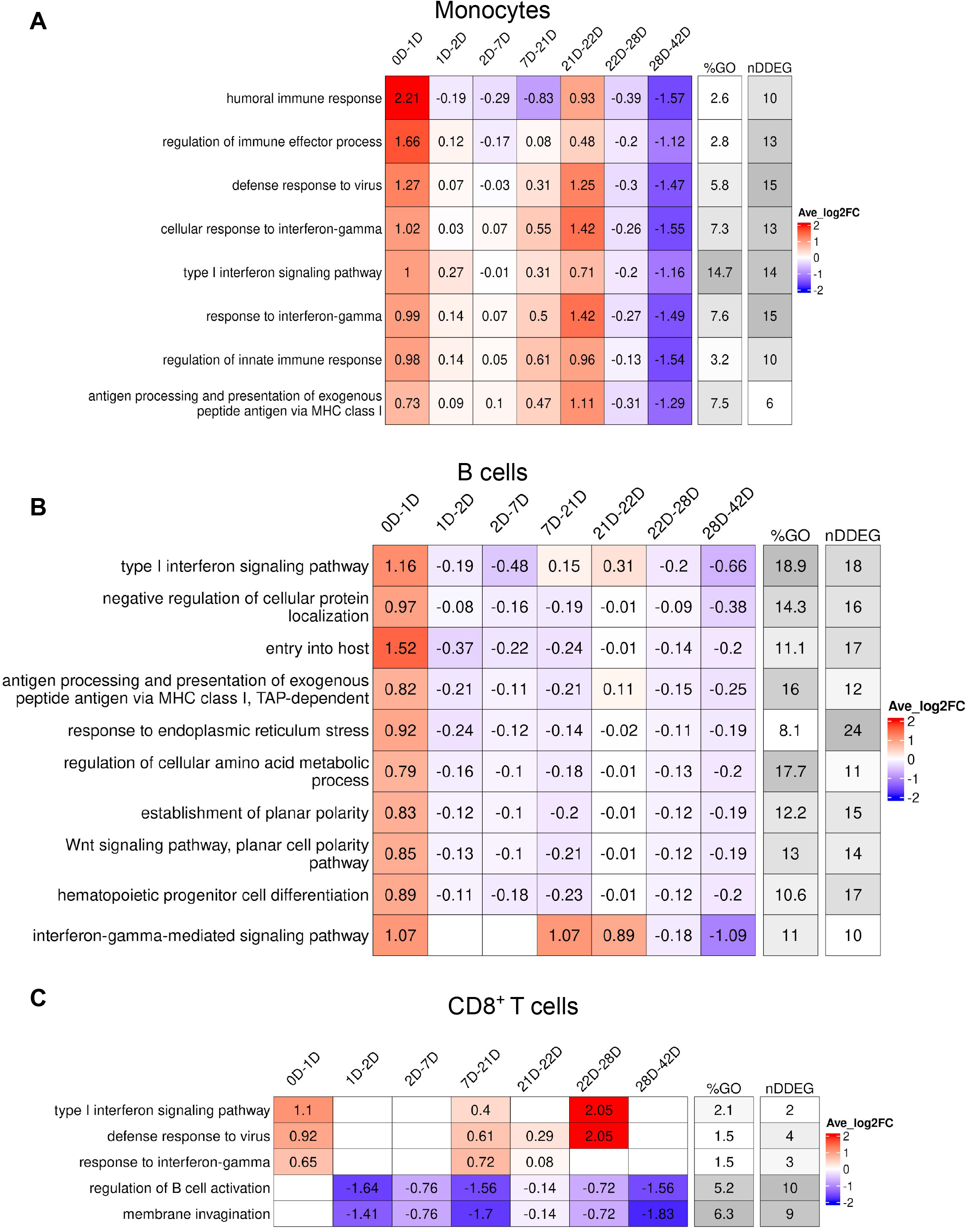
TimeHeatmap of monocytes, B cells and CD8^+^ T cells. (A) *TimeHeatmap* of monocytes. (B) *TimeHeatmap* of B cells. (C) *TimeHeatmap* of CD8^+^T cells. Day 1 is the first dose of vaccination, Day 21 is the second dose of vaccination.

## References

1. Bar-Joseph Z, et al. Studying and modelling dynamic biological processes using time-series gene expression data. Nat Rev Genet. 2012;13(8):552–64.

2. Hwang B, et al. Single-cell RNA sequencing technologies and bioinformatics pipelines. Experimental and Molecular Medicine. 2018;50.

3. Love MI, et al. Moderated estimation of fold change and dispersion for RNA-seq data with DESeq2. Genome Biol. 2014;15(12):550.

4. McCarthy DJ, et al. Differential expression analysis of multifactor RNA-Seq experiments with respect to biological variation. Nucleic Acids Res. 2012;40(10):4288–97.

5. Ritchie ME, et al. limma powers differential expression analyses for RNA-sequencing and microarray studies. Nucleic Acids Res. 2015;43(7):e47.

6. Fischer DS, et al. Impulse model-based differential expression analysis of time course sequencing data. Nucleic Acids Res. 2018;46(20):e119.

7. Nueda MJ, et al. Next maSigPro: updating maSigPro bioconductor package for RNA-seq time series. Bioinformatics. 2014;30(18):2598–602.

8. Zhou F, et al. Clinical course and risk factors for mortality of adult inpatients with COVID- 19 in Wuhan, China: a retrospective cohort study. Lancet. 2020;395(10229):1054–62.

9. Wu ZY, and McGoogan JM. Characteristics of and Important Lessons From the Coronavirus Disease 2019 (COVID-19) Outbreak in China Summary of a Report of 72 314 Cases From the Chinese Center for Disease Control and Prevention. Jama-Journal of the American Medical Association. 2020;323(13):1239–42.

10. Pedersen SF, and Ho YC. SARS-CoV-2: a storm is raging. Journal of Clinical Investigation. 2020;130(5):2202–5.

11. Bergamaschi L, et al. Longitudinal analysis reveals that delayed bystander CD8(+) T cell activation and early immune pathology distinguish severe COVID-19 from mild disease. Immunity. 2021;54(6):1257-+.

12. Bernardes JP, et al. Longitudinal Multi-omics Analyses Identify Responses of Megakaryocytes, Erythroid Cells, and Plasmablasts as Hallmarks of Severe COVID-19. Immunity. 2020;53(6):1296-+.

13. Aid M, et al. Vascular Disease and Thrombosis in SARS-CoV-2-Infected Rhesus Macaques. Cell. 2020;183(5):1354-+.

14. Zhu LN, et al. Single-Cell Sequencing of Peripheral Mononuclear Cells Reveals Distinct Immune Response Landscapes of COVID-19 and Influenza Patients. Immunity. 2020;53(3):685-+.

15. Liu C, et al. Time-resolved systems immunology reveals a late juncture linked to fatal COVID-19. Cell. 2021;184(7):1836-+.

16. Arunachalam PS, et al. Systems vaccinology of the BNT162b2 mRNA vaccine in humans. Nature. 2021;596(7872):410-+.

17. Hao YH, et al. Integrated analysis of multimodal single-cell data. Cell. 2021;184(13):3573-+.

18. Aran D, et al. Reference-based analysis of lung single-cell sequencing reveals a transitional profibrotic macrophage. Nature Immunology. 2019;20(2):163-+.

19. Krieger G, et al. Independent evolution of transcript abundance and gene regulatory dynamics. Genome Research. 2020;30(7):1000–11.

20. Yuan Y, and Bar-Joseph Z. Deep learning for inferring gene relationships from single-cell expression data. Proceedings of the National Academy of Sciences of the United States of America. 2019;116(52):27151–8.

21. Graw F. DISEASE Deciphering the triad of infection, immunity and pathology. Elife. 2021;10.

22. Van den Berge K, et al. Trajectory-based differential expression analysis for single-cell sequencing data. Nature Communications. 2020;11(1).

23. Qiu XJ, et al. Reversed graph embedding resolves complex single-cell trajectories. Nature Methods. 2017;14(10):979-+.

24. Chen G, et al. Clinical and immunological features of severe and moderate coronavirus disease 2019. Journal of Clinical Investigation. 2020;130(5):2620–9.

25. Gong J, et al. Correlation analysis between disease severity and inflammation-related parameters in patients with COVID-19: a retrospective study. Bmc Infectious Diseases. 2020;20(1).

26. Mehta P, et al. COVID-19: consider cytokine storm syndromes and immunosuppression. Lancet. 2020;395(10229):1033–4.

27. Qin C, et al. Dysregulation of Immune Response in Patients With Coronavirus 2019 (COVID-19) in Wuhan, China. Clinical Infectious Diseases. 2020;71(15):762–8.

28. Goyal A, et al. Potency and timing of antiviral therapy as determinants of duration of SARS-CoV-2 shedding and intensity of inflammatory response. Science Advances. 2020;6(47).

29. Ackermann M, et al. Patients with COVID-19: in the dark-NETs of neutrophils. Cell Death and Differentiation. 2021;28(11):3125–39.

30. Morrissey SM, et al. A specific low-density neutrophil population correlates with hypercoagulation and disease severity in hospitalized COVID-19 patients. Jci Insight. 2021;6(9).

31. Cambier S, et al. Atypical response to bacterial co-infection and persistent neutrophilic broncho-alveolar inflammation distinguish critical COVID-19 from influenza. JCI Insight. 2021.

32. Kaiser R, et al. Self-sustaining IL-8 loops drive a prothrombotic neutrophil phenotype in severe COVID-19. Jci Insight. 2021;6(18).

33. Freire PP, et al. The relationship between cytokine and neutrophil gene network distinguishes SARS-CoV-2-infected patients by sex and age. Jci Insight. 2021;6(10).

34. Libby P, and Luscher T. COVID-19 is, in the end, an endothelial disease. European Heart Journal. 2020;41(32):3038–44.

35. Bardoel BW, et al. The Balancing Act of Neutrophils. Cell Host & Microbe. 2014;15(5):526–36.

36. Metzemaekers M, et al. Kinetics of peripheral blood neutrophils in severe coronavirus disease 2019. Clinical & Translational Immunology. 2021;10(4).

37. Papayannopoulos V. Neutrophil extracellular traps in immunity and disease. Nature Reviews Immunology. 2018;18(2):134–47.

38. Zuo Y, et al. Autoantibodies stabilize neutrophil extracellular traps in COVID-19. Jci Insight. 2021;6(15).

39. Zuo Y, et al. Neutrophil extracellular traps in COVID-19. JCI Insight. 2020;5(11).

40. Merad M, and Martin JC. Pathological inflammation in patients with COVID-19: a key role for monocytes and macrophages. Nature Reviews Immunology. 2020;20(6):355–62.

41. Grant RA, et al. Circuits between infected macrophages and T cells in SARS-CoV-2 pneumonia. Nature. 2021;590(7847).

42. Ackermann M, et al. Pulmonary Vascular Endothelialitis, Thrombosis, and Angiogenesis in Covid-19. New England Journal of Medicine. 2020;383(2):120–8.

43. Rapkiewicz AV, et al. Megakaryocytes and platelet-fibrin thrombi characterize multi-organ thrombosis at autopsy in COVID-19: A case series. Eclinicalmedicine. 2020;24.

44. Schmaier AA, et al. Tie2 activation protects against prothrombotic endothelial dysfunction in COVID-19. Jci Insight. 2021;6(20).

45. Del Valle DM, et al. An inflammatory cytokine signature predicts COVID-19 severity and survival. Nature Medicine. 2020;26(10):1636-+.

46. Leisman DE, et al. Cytokine elevation in severe and critical COVID-19: a rapid systematic review, meta-analysis, and comparison with other inflammatory syndromes. Lancet Respiratory Medicine. 2020;8(12):1233–44.

47. Skendros P, et al. Complement and tissue factor-enriched neutrophil extracellular traps are key drivers in COVID-19 immunothrombosis. Journal of Clinical Investigation. 2020;130(11):6151–7.

48. Carfi A, et al. Persistent Symptoms in Patients After Acute COVID-19. Jama-Journal of the American Medical Association. 2020;324(6):603–5.

49. Huang CL, et al. 6-month consequences of COVID-19 in patients discharged from hospital: a cohort study. Lancet. 2021;397(10270):220–32.

50. Nalbandian A, et al. Post-acute COVID-19 syndrome. Nature Medicine. 2021;27(4):601–15.

51. McNab F, et al. Type I interferons in infectious disease. Nature Reviews Immunology. 2015;15(2):87–103.

52. Muller U, et al. FUNCTIONAL-ROLE OF TYPE-I AND TYPE-II INTERFERONS IN ANTIVIRAL DEFENSE. Science. 1994;264(5167):1918-21.

53. Blanco-Melo D, et al. Imbalanced Host Response to SARS-CoV-2 Drives Development of COVID-19. Cell. 2020;181(5):1036-+.

54. Hadjadj J, et al. Impaired type I interferon activity and inflammatory responses in severe COVID-19 patients. Science. 2020;369(6504):718-+.

55. Darazam IA, et al. Role of interferon therapy in severe COVID-19: the COVIFERON randomized controlled trial. Scientific Reports. 2021;11(1).

56. Loftus RM, and Finlay DK. Immunometabolism: Cellular Metabolism Turns Immune Regulator. Journal of Biological Chemistry. 2016;291(1):1–10.

57. O’Neill LAJ, et al. A guide to immunometabolism for immunologists. Nature Reviews Immunology. 2016;16(9):553–65.

58. O’Brien KL, and Finlay DK. Immunometabolism and natural killer cell responses. Nature Reviews Immunology. 2019;19(5):282–90.

59. C G. Smoothing spline ANOVA models: R package gss. J Stat Softw. 2014;58(5):25.

60. Metwally AA, et al. MetaLonDA: a flexible R package for identifying time intervals of differentially abundant features in metagenomic longitudinal studies. Microbiome. 2018;6(1):32.

61. Yu G, et al. clusterProfiler: an R package for comparing biological themes among gene clusters. OMICS. 2012;16(5):284–7.

62. Kuleshov MV, et al. Enrichr: a comprehensive gene set enrichment analysis web server 2016 update. Nucleic Acids Res. 2016;44(W1):W90–7.

63. Gu ZG, et al. Complex heatmaps reveal patterns and correlations in multidimensional genomic data. Bioinformatics. 2016;32(18):2847–9.

